# Can we improve slow learning in cerebellar patients?

**DOI:** 10.1101/2020.07.03.185959

**Authors:** Thomas Hulst, Ariels Mamlins, Maarten Frens, Dae-In Chang, Sophia L. Göricke, Dagmar Timmann, Opher Donchin

## Abstract

Cerebellar patients are impaired in motor adaptation. Further, motor adaptation can be divided into a fast component and a slow component, and the cerebellum is known to be especially crucial for the slow component. We tested whether the cerebellar deficit in motor adaptation can be ameliorated by training paradigms targeting slow learning using four visuomotor tasks: standard, gradual, overlearning and long intertrial intervals. We measured slow learning in patients and age-matched controls using a standard paradigm developed for reaching movements by Smith in 2006. The paradigm quantifies slow learning as the magnitude of spontaneous recovery of a previously learned and washed-out adaptation. Cerebellar patients had slower learning and reached a lower level of final adaptation, as seen in previous studies. Nevertheless, both groups had robust spontaneous recovery. Moreover, spontaneous recovery was increased in both groups in the overlearning paradigm with no significant difference between them. Computational modeling suggested that increased spontaneous recovery in the Control group reflects a changed slow adaptation system. That is, in the overlearning in controls, the slow system forgot more slowly and had a slower response to errors. In contrast, the same modeling suggests that no such change in the slow system occurred in the Cerebellar group. Rather, increased spontaneous recovery in this group seems to be the result of greater accumulated slow adaptation in the overlearning. We used our modeling results to predict that overlearning should have slower adaptation in the counterperturbation. Modeling further suggested we would not see these difference in the cerebellar patients. Follow-up analysis confirmed these model predictions. Taken together, our results imply that residual slow learning in cerebellar patients is expressed during increased training trials, but the primary cerebellar deficit is not improved.

## 1 Introduction

Patients with cerebellar degeneration exhibit a range of impairments in motor control, including incoordination of eye movements, dysarthria, limb incoordination, and gait disturbances (Mariotti et al., 2005), as well as impairments in the cognitive domain (Schmahmann and Sherman, 1998). Rehabilitation, including physical therapy, speech therapy and occupational therapy, is a primary approach to improve quality of life for cerebellar patients, and the consensus is that cerebellar patients benefit from rehabilitation. However, there are many open questions about how to maximize efficacy. For instance, a challenge in effective rehabilitation is that cerebellar patients suffer from motor learning deficits (Maschke et al., 2004; Tseng et al., 2007) and the motor learning deficits affect the success of neurorehabilitation programs (Hatakenaka et al., 2012). One important question is whether specific training protocols can alleviate motor learning deficits. We address the question of whether specific training can change motor learning deficits in cerebellar patients in the context of reach adaptation.

Much research on motor learning has focused on reach adaptation, leading to a well-characterized task that serves as good model for more general questions in motor learning and adaptation (Krakauer et al., 2019; Wang et al., 2024). In reach adaptation, movements of the arm towards a target are perturbed such that movement of the cursor deviates from actual movement of the hand. This leads to corrections on subsequent movements. Within these corrections, two learning processes can be distinguished: fast -- thought to reflect explicit strategic and reinforcement learning; and slow -- thought to reflect implicit sensory error-based learning (McDougle et al., 2015; Smith et al., 2006). Although there is some indication that the cerebellum contributes to both processes (Butcher et al., 2017; Therrien et al., 2016), there is extensive evidence specifically for a deficit in the slow process (Izawa and Shadmehr, 2011; Morehead et al., 2017; Tseng et al., 2007; among others).

The question of whether training paradigms specifically designed to create more slow learning can ameliorate adaptation deficits in cerebellar patients remains open. Early evidence indicated beneficial effects in gradual introduction of reaching movement perturbations in cerebellar patients (Criscimagna-Hemminger et al., 2010), but the results were not replicated in subsequent work (Gibo et al., 2013; Schlerf et al., 2013). Other paradigms shown to emphasize slow learning in healthy subjects have not been explored in cerebellar patients. We consider two additional such paradigms. The first is overlearning, continued training after asymptotic performance, which increases retention in healthy subjects as a function of the number of trials trained at asymptote (Joiner and Smith, 2008). The second uses long intertrial intervals (ITI) between movements. This decreases the rate of learning in healthy subjects, but increases retention (Kim et al., 2015; Sing et al., 2009).

The aim of the present study is to test all the paradigms in the same sets of healthy subjects and cerebellar patients. We tested twenty patients with degenerative ataxia and twenty healthy age- and sex-matched controls on a visuomotor reaching adaptation task under four different training paradigms. We tested their effects on the development of slow learning using a behavioral measure called spontaneous recovery. In spontaneous recovery, we measure the tendency of subjects to return to a learned perturbation after application of a short counterperturbation. Spontaneous recovery is thought to reflect retained slow learning following wash out of the fast component (Coltman et al., 2019; Smith et al., 2006).

In addition to measuring spontaneous recovery, we also characterized the extent of fast and slow adaptation using a two-state model based on (Smith et al., 2006)). Although this model has known limitations (Forano and Franklin, 2020; Petitet et al., 2018; Zarahn et al., 2008), it provides good fit to human behavior in many paradigms and successfully isolates the two main time constants of human learning. By using the model to characterize slow learning, we can gain insight into the changes caused by the behavioral paradigms.

Because this research is the first to compare multiple paradigms thought to increase slow learning, our interest was both in the extent to which slow learning increased in healthy subjects and in the difference between healthy subjects and cerebellar patients. If there is a way to increase the amount of slow learning in cerebellar patients, this could be used to better optimize training schedules in rehabilitation.

## 2 Methods

### 2.1 Participants

Twenty participants with cerebellar degeneration (9 females, 54.9 years ± 10.8 (SD), range 18 – 70 years) and twenty age- and sex-matched participants (9 females, 55.2 years ± 11.2 (SD), range 18 – 71 years) took part in the study. Cerebellar participants were recruited from the patients attending our ataxia clinic and matched controls were recruited via print advertisements distributed on the hospital campus. Only right-handed individuals were included, as assessed by the Edinburgh Handedness Inventory (Oldfield, 1971). The severity of cerebellar symptoms in the group of cerebellar participants was assessed by one experienced neurologist (DT) and healthy age- and sex-matched controls were examined by AM. Cerebellar symptoms were scored on the International Cooperative Ataxia Rating Scale (ICARS; Trouillas et al., 1997), as well as the Scale for the Assessment and Rating of Ataxia (SARA; Schmitz-Hübsch et al., 2006). The group of cerebellar participants was diagnosed with diseases known to primarily affect the cerebellar cortex (Gomez et al., 1997; Timmann et al., 2009). Three age-matched controls were excluded and replaced due to neurological symptoms on their examination or minor extracerebellar pathology on their MRI. All participants gave informed oral and written consent. The experiment was approved by the ethics committee of the medical faculty of the University of Duisburg-Essen and conducted in accordance with the Declaration of Helsinki. The characteristics of the recruited cerebellar participants and matched controls can be found in **Table 1**.

**Table 1.** Overview Cerebellar participants and Control participants.

Cerebellar participants were age- and sex-matched with control participants on the right side of the table.
| Cerebellar participants |  |  |  |  |  |  | Control participants |  |  |
| --- | --- | --- | --- | --- | --- | --- | --- | --- | --- |
| ID | Age | Sex | Diagnosis | Disease duration | ICARS (total/100) | SARA (total/40) | ID | Age | Sex |
| P01 | 18 | M | ADCAIII | 18 years | 10.5 | 5 | C01 | 18 | M |
| P02 | 47 | M | ADCAIII | 25+ years | 40 | 17.5 | C02 | 43 | M |
| P03 | 50 | F | ADCAIII | 25+ years | 31 | 11.5 | C03 | 50 | F |
| P04 | 51 | M | SCA14 | 25+ years | 31 | 11.5 | C04 | 50 | M |
| P05 | 51 | F | ADCAIII | 19 years | 29 | 12.5 | C05 | 51 | F |
| P06 | 52 | F | SCA14 | 22 years | 26.5 | 12 | C06 | 53 | F |
| P07 | 53 | F | SCA6 | 2 years | 23 | 9.5 | C07 | 54 | F |
| P08 | 53 | M | SAOA | 17 years | 40 | 15 | C08 | 53 | M |
| P09 | 53 | M | ADCAIII | 17 years | 36 | 11 | C09 | 51 | M |
| P10 | 54 | F | SCA6 | 4 years | 30.5 | 10 | C10 | 56 | F |
| P11 | 56 | F | SCA14 | 25+ years | 28 | 12 | C11 | 58 | F |
| P12 | 57 | M | SCA6 | 11 years | 38 | 11 | C12 | 53 | M |
| P13 | 57 | M | SCA6 | 15 years | 28 | 8 | C13 | 59 | M |
| P14 | 58 | M | SAOA | 25+ years | 63.5 | 22 | C14 | 61 | M |
| P15 | 59 | M | SCA6 | 5 years | 23 | 9.5 | C15 | 63 | M |
| P16 | 60 | F | SCA6 | 11 years | 36.5 | 14 | C16 | 60 | F |
| P17 | 63 | F | ADCAIII | 23 years | 33 | 13.5 | C17 | 66 | F |
| P18 | 66 | M | SAOA | 13 years | 24.5 | 11 | C18 | 66 | M |
| P19 | 70 | F | SAOA | 7 years | 32.5 | 12.5 | C19 | 71 | F |
| P20 | 70 | M | SAOA | 16 years | 38 | 15 | C20 | 67 | M |

### 2.2 Task

All participants completed a standard reaching task with visuomotor perturbations. The experimental setup and task have been described previously in other studies from our group (Rabe et al., 2009). In short, participants were seated in front of an upright monitor and could, with their right hand, move a two-jointed manipulandum freely in the horizontal plane (**Figure 1A**). Vision of the participant’s arm was obstructed by a black cloth. Hand position and velocities were measured in a resolution of 10^6^ counts per revolution and a sampling rate of 200 Hz (DMC-1826; Galil Motion Control). The location of the participant’s hand was represented on the monitor by a green dot with a diameter of 5 mm. The origin and target locations were represented by a circle with a diameter of 10 mm, colored red and white respectively. At the start of each trial, the participant’s hand was moved towards the origin location by the servomotors connected to the manipulandum. Then, after a delay of 2 sec, a target circle appeared at one of three possible target locations, located 10 centimeters away from the origin at an angle of 66°, 90° or 114° (**Figure 1B**). Participants were instructed to move the green dot from the origin towards the target with a “quick and accurate movement” as soon as the target appeared. When participants moved the cursor through an invisible boundary located 10 centimeters from the origin, their hand was gently brought to a stop by a simulated cushion, indicating the end of the movement. Following each movement, participants received feedback on whether they hit the target and moved with the correct velocity. The target turned yellow when moving too fast, blue when moving too slow, and green when moving with the correct velocity. Participants moved with the correct velocity when their movement and reaction time fell within a 250 ms window centered around 500 ms. The 250 ms window shrunk by 10% every time a movement had the correct velocity and increased by 10% when moving too fast or slow, adapting to a participant’s individual capabilities. When participants also managed to hit the target, in addition to moving with the correct velocity, a “yahoo!” sound was played.

**Figure 1:**
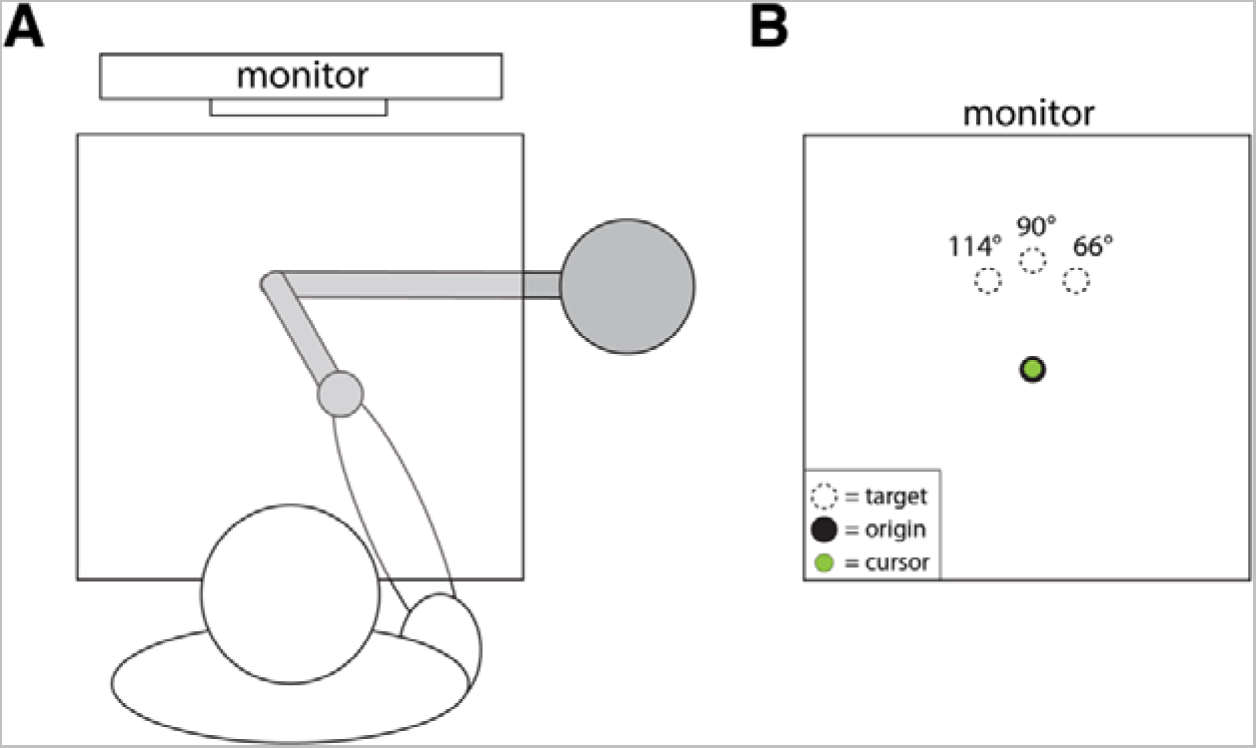
A) Experimental setup. Subject grips a two-dimensional manipulandum that controls movements of a cursor on an upright monitor which is used to show targets as well. An opaque tabletop obstructed the participant’s view of their hand and the robot arm. Additionally, a black cloth was draped over the shoulders of the participant and attached to the table to obstruct vision of the arm. B) Location of the origin and targets on the monitor. On each trial, participants start by moving their cursor to the origin. After a 2 sec interval, one of the three targets would be shown. Participants were instructed to move their cursor “as quickly and as accurately” to the target.

All participants completed four training paradigms (standard, overtraining, gradual, and long intertrial interval), with the order of paradigms was counterbalanced with a Latin-squares design against first-order carryover effects (Williams, 1949). Every paradigm order was completed by 10 participants each (five cerebellar participants and five control participants, **Figure 2A**). Analysis of the effects of counterbalancing are presented in the supplementary materials. Participants were allowed to take 5- to 10-minute breaks between paradigms.

**Figure 2:**
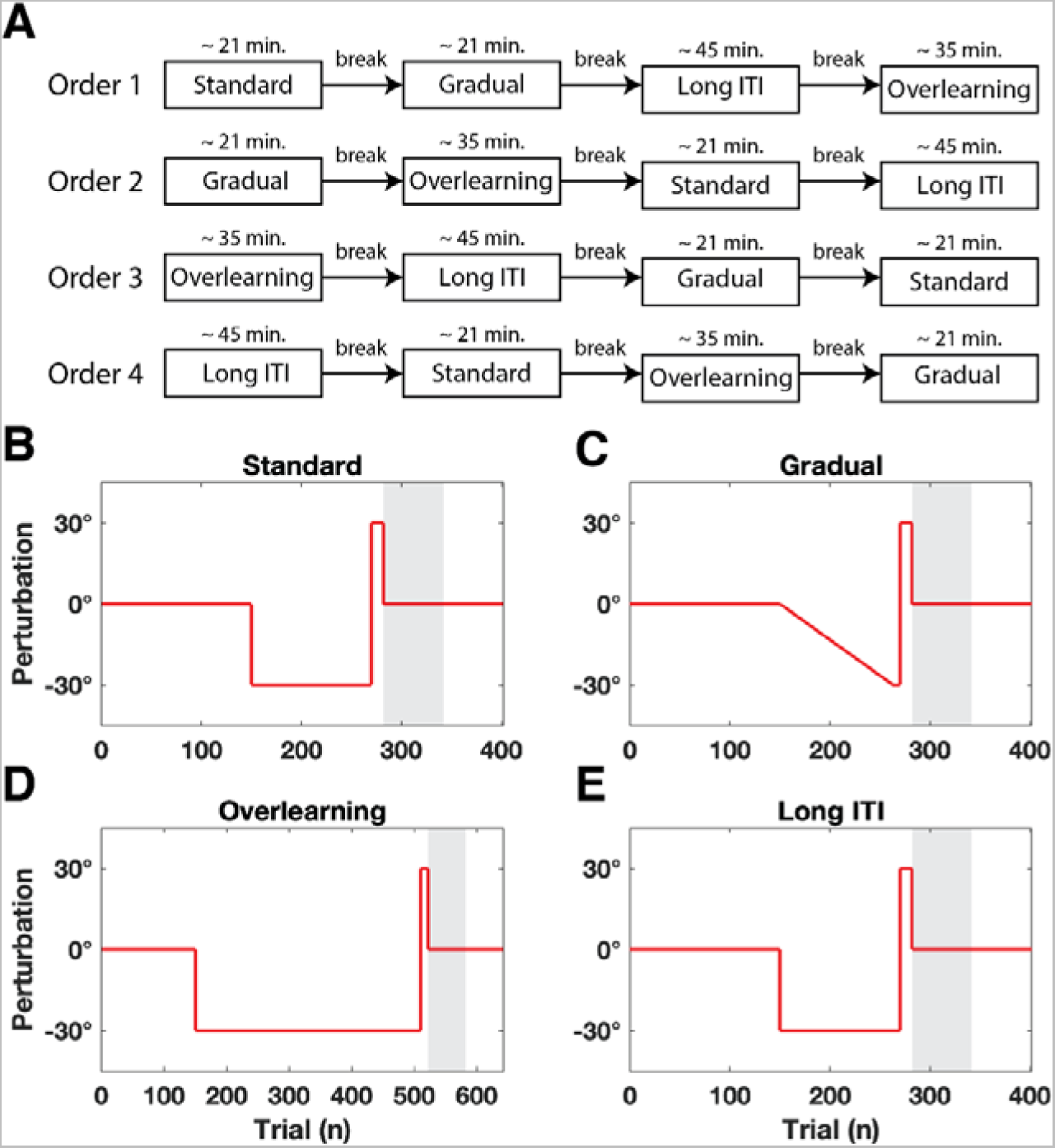
A) Overview of the paradigm order counterbalanced using a Latin Square. B–E) Trial structure of the experimental paradigms. Red line indicates the size and direction of the visuomotor perturbation. Perturbation is always shown as clockwise although it actually alternated on successive sets for each subject. Grey area indicates the block of 60 clamp trials where the cursor moves straight to the target regardless of hand movement. Clamp trials were also pseudo-randomly interspersed during baseline and adaptation sets (not pictured).

Each paradigm consisted of three sets: a baseline set, an adaptation set, and a washout set. The baseline set consisted of 135 null trials, in which participants received veridical feedback on hand position, and 15 pseudo-randomly interspersed clamp trials. Clamp trials are trials in which the cursor moved directly to the target regardless of actual hand movement.

The adaptation set introduced adaptation trials with a 30° rotation of the cursor movement relative to the hand. This rotation was in a different direction for successive sets for each subject: clockwise in their first set, counterclockwise in their second set, and so on. The adaptation sets were different in each paradigm:

*Standard paradigm*: the adaptation set was 108 adaptation trials with 12 pseudo-randomly interspersed clamp trials (**Figure 2B**).

*Gradual paradigm*: the adaptation set was 108 adaptation trials where the visuomotor perturbation was increased linearly over the course of the set. The final 6 trials were full adaptation trials. In addition, 12 clamp trials were pseudo-randomly interspersed (**Figure 2C**). *Overlearning paradigm:* the adaptation set was 324 adaptation trials and 36 interspersed clamp trials (**Figure 2D**).

*ITI paradigm:* the adaptation set was 120 adaptation trials but the delay in the origin was 15 seconds instead of 2 seconds (**Figure 2E**). The adaptation set of the long ITI paradigm did not include any clamp trials, thus trial-to-trial forgetting was only dependent on the passage of time.

The adaptation set was followed by the washout set. The first 12 trials of the washout set where counterperturbation trials, where the direction of the perturbation was flipped from the direction in the adaptation set. These were followed by 60 clamp trials and 60 null trials.

### 2.3 MR imaging

Cerebellar participants and age-matched controls were examined in a 3T combined MRI-PET system (Siemens Healthcare, Erlangen, Germany) with a 16-channel head coil (Siemens Healthcare) [TR = 2,530 ms; TE = 3.26 ms, TI = 1,100 ms; flip angle 7 deg; voxel size 0.5 × 0.5 × 1.0 mm³]. All MR scans were evaluated by an experienced neuroradiologist (SLG).

A voxel-based morphometry analysis was applied to the cerebellum of each participant as described previously (Hulst et al., 2015; Taig et al., 2012). The analysis was automated with an in-house program written for MATLAB 9.4 using the SUIT toolbox (version 3.2) (Diedrichsen et al., 2009), implemented in SPM12 (http://www.fil.ion.ucl.ac.uk/spm/software/spm12). A resampling procedure (permutation test) was conducted to control the family-wise error rate. Using this procedure, we found that under arbitrary divisions of the subjects into two groups, there was less than a 5% chance of finding any significant voxels by chance if we set the significance threshold at an absolute t-score of 3.95. This is what we did.

### 2.4 Analysis of behavioral data

Behavioral data was analyzed in MATLAB 9.11 (MathWorks, Natick, USA). Our primary outcome measure was the reaching direction (in degrees) at the end of the movement (i.e., when participants hit the simulated cushion). Since the cushion allowed movements to be ballistic, they were straight, and this outcome measure was highly correlated with any other that is commonly used such as hand angle at maximum velocity or at a specific time in the movement. This was tested by redoing the analysis with two other outcome measures without significantly changing the results.

The reaching direction was calculated by taking the angle between a straight line from the position of movement onset to the target and a straight line from the position of movement onset to hand position at the end of the movement. Movement onset was defined as the first moment when movement velocity exceeded 5cm/s. Reaching directions were corrected for movement biases by calculating the average reaching direction in each baseline set and subtracting this from the subsequent adaptation and washout sets of a training paradigm. For ease of interpretation, reaching directions were flipped towards the same direction, regardless of perturbation direction, in all figures and analyses. Furthermore, paradigms were reordered to a canonical order for each participant, starting with the standard learning paradigm, then gradual learning, overlearning and finally the long ITI paradigm, regardless of the order the participant encountered the paradigms.

Statistical analyses were conducted using Markov Chain Monte Carlo (MCMC) methods in MATLAB and JAGS 4.3.0 (Plummer, 2003). A mixed-design model (ANOVA-like) was used to estimate the difference in reaching directions between factors. Participant group (cerebellar participant or control participant) was included as a between-subject factor. Movement phase and training paradigm were included as within-subject factors. Movement phases were defined as follows: baseline (all 150 trials in the baseline set), early adaptation (first 30 trials of the adaptation set), late adaptation (final 6 trials of the adaptation set) and recovery (all 60 clamp trials in the washout set). A random intercept for each participant and phase was included as well. The model ran on four separate chains with an adaptation phase of 2,500 samples and burn-in phase of 12,500 samples, after which we collected 25,000 samples per chain. The MCMC procedure gives us a posterior distribution of credible parameter values, given the data. For point estimates of parameters we used the maximum (modal) value of this posterior probability distribution. Uncertainty in the point estimate was expressed using the 95% highest density interval (HDI). The 95% HDI contains 95% of the mass of credible parameter values, where each value within the HDI has a higher probability density than any value outside the HDI. When the HDI falls completely within the Region of Practical Equivalence (ROPE), we accept the null value of the parameter and when the HDI falls completely outside the ROPE, we reject the null value of the parameter. The ROPE was set at [−2°, 2°] to match the variability of the mean of the baseline across paradigms. Each parameter was visually and quantitatively checked to assure proper sampling of the posterior distribution using common MCMC diagnostics (Kruschke, 2010). First, trace plots were visually inspected for chain convergence. Next, the effective sample size (ESS), the potential scale reduction factor (PSRF) and the Monte Carlo Standard Error (MCSE) were calculated for all parameters. The PSRF was close to 1 for each parameter (max: 1.0002), MCSE was close to 0 for each parameter (max: 0.0001), and median ESS was generally large (>> 5,000), indicating convergence of the model run. The model code is available on https://github.com/Motor-Learning-Lab/paper-slowlearning.

Our within-subject design might lead to order effects. That is the particular order in which a subject did the paradigms might affect their performance on each paradigm. As mentioned above, order was counterbalanced between subjects using a latin squares approach and this should largely eliminate order effects. Order effects are not thought to affect the slow system (Avraham et al., 2021; Morehead et al., 2015) and are thought to be prevented by immediate washout followed by a delay (Kitago et al., 2013). Nevertheless, to further address this concern, we tested for meaningful order effects and for the possibility that order effects might explain our other results. To do this, we fit two models that included order effects to the data. Methods and results for this analysis are included in the supplementary materials.

### 2.5 State-space modeling

A two-state model of motor learning was fit to the reaching directions of all trials in each individual participant. Our model was based on that originally presented by Smith and colleagues (Smith et al., 2006) and posits a fast state (x^F^), that learns and forgets quickly (the parameters of learning and forgetting are called BFast and AFast, respectively), and a slow state (x ^S^), that learns and forgets slowly (with parameters BSlow and ASlow). Adaptation in both states is driven by the mismatch between visual cursor and target. The model was fit using a Bayesian approach similar to that in van der Vliet et al. (2018). Details of the model and the Bayesian fitting procedure used to estimate individual subject parameters are available in the supplementary materials. The model code is available on https://github.com/Motor-Learning-Lab/paper-slowlearning.

The ability of the model to capture the key features of the data was assessed using posterior predictive plots. Posterior predictive plots show the distribution of data that can be generated by the model given the posterior distribution of the parameters. That is, for each subject we can generate a new set of data that the model sees as similar to the data from that subject. For the posterior predictive plots in this paper, we produced 1,000 datasets for each subject and from this produced 1,000 estimated mean datasets. We used this to get point estimates of the average adaptation level as well as the levels of both slow and fast learning. We also generated 95% HDIs for each of these quantities.

As can be seen in the results, our central result is an increased spontaneous recovery in the overlearning paradigm. The goal of modeling was to better understand this increase. To this purpose, we compared the estimated learning and forgetting rates of the fast system (BFast and AFast, respectively) and slow system (BSlow and ASlow) across paradigms and groups, both at the level of individual subjects and the population. The paper focuses on the comparison of the standard and overlearning paradigms, and full results are given in the supplementary material.

### 2.6 Post-hoc model validation

As made clear in the Results section, the results from the model suggested that overlearning in controls may have led to more stability in the slow state: less learning combined with less forgetting when compared to the standard paradigm. We did not see the same change in the parameters of the cerebellar participants between the overlearning and standard paradigms.

This led us to predict that rate of adaptation would be affected by overlearning. To test this, we compared the learning rate before and after the adaptation set. That is, we compared rate in initial adaptation (before the overlearning) to learning rate in the counterperturbation (after the overlearning). We expected the learning rate to be slower in the overlearning sets for controls and the same for cerebellar participants. We expected there to be no difference in the standard set. To do this comparison, we used the same hierarchical Bayesian approach as before to find the slope in the first 13 trials of the adaptation and counterperturbation phases.

## 3 Results

### 3.1 Voxel-based morphometry (VBM)

First, we confirmed that our patient group has prominent cerebellar degeneration compared to the matched control group with structural MRI (VBM). **Figure 3** displays the difference in gray matter volume per voxel (in t-scores) between healthy participants and cerebellar participants. Using the significance threshold set based on our permutation analysis, no significant positive differences were found; thus, the figure displays negative t-scores only.

**Figure 3:**
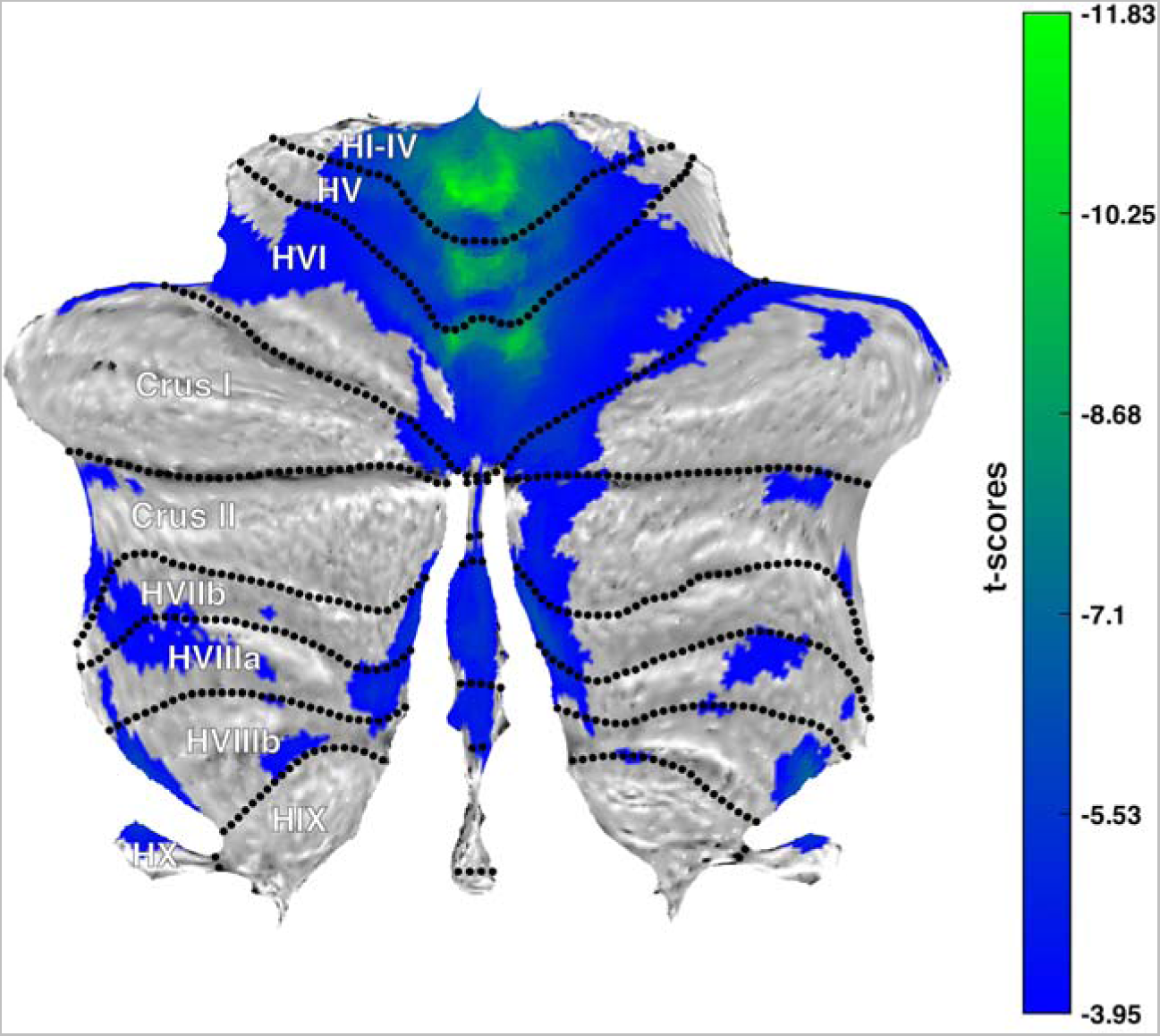
Flatmap of the cerebellum. Colors indicate the gray matter volume difference per voxel between healthy participants and cerebellar participants in t-scores. Voxels that do not exceed the threshold (−3.95) are not colored, lower significant t-scores are colored blue, and higher significant t-scores are colored green. Flatmap template from Diedrichsen and Zotow, 2015.

The VBM analysis revealed a pattern of cerebellar degeneration in patients largely consistent with prior work (Hulst et al., 2015). The volume loss was largest in the anterior lobe of the cerebellum and the superior part of the posterior lobe (i.e., lobule VI). Cerebellar degeneration of the anterior cerebellum and lobule VI (i.e. the anterior hand area) are associated with motor learning deficits (Donchin et al., 2012; Rabe et al., 2009). Cerebellar degeneration was less pronounced in the inferior parts of the posterior lobe compared to earlier work (Hulst et al., 2015), which could be explained by younger cerebellar participants in the current study, with less severe ataxia scores.

### 3.2 Average reaching directions

The average reaching directions for each paradigm in control participants and cerebellar participants are plotted in **Figure 4A** and **4B**. Differences in mean reaching directions per phase, paradigm and subject group were tested using the mixed-model design from the methods section and the most important results are described below.

**Figure 4:**
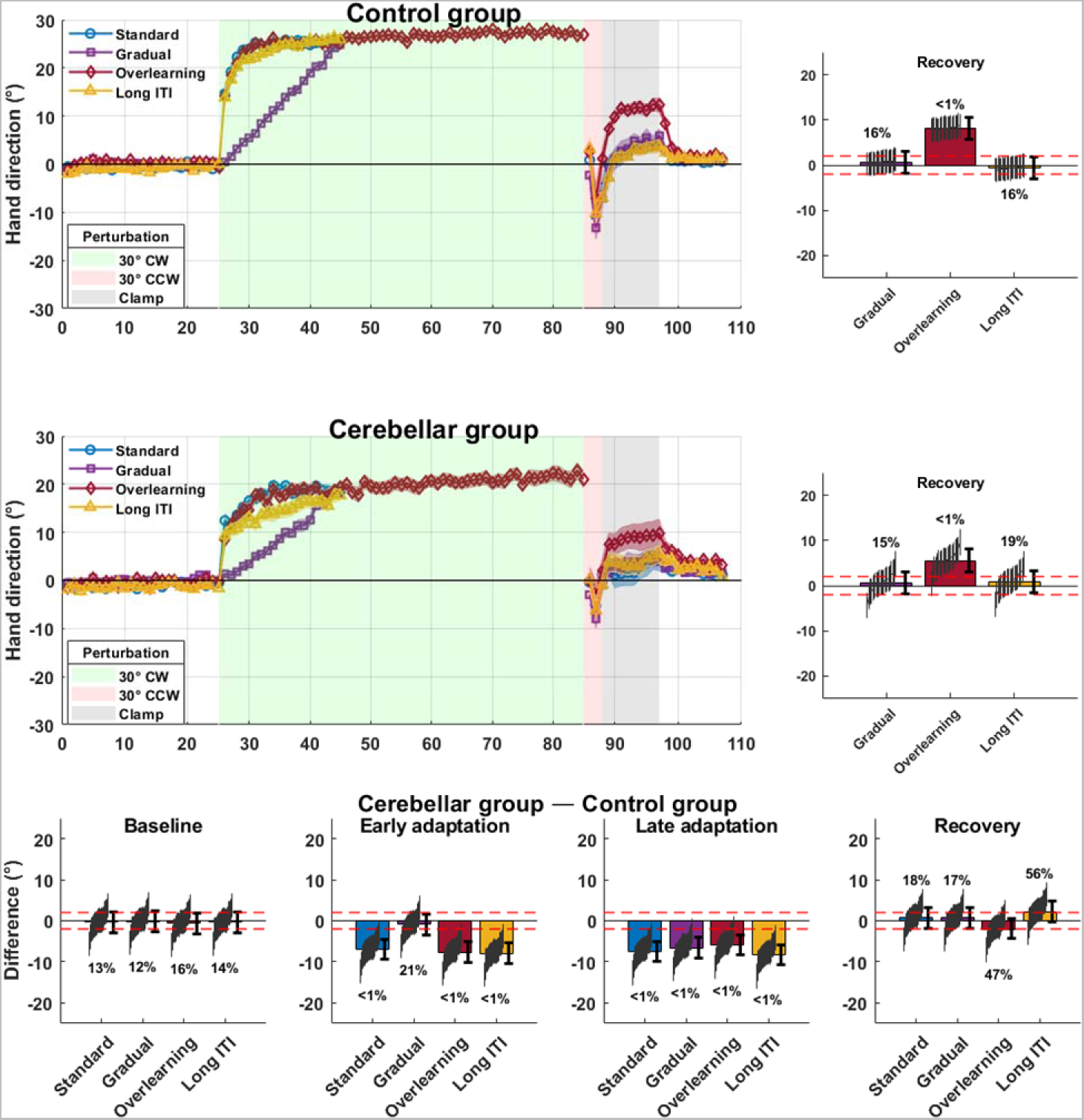
Average reaching directions of control participants and cerebellar participants. A) Reaching directions of control participants. Trials were binned per 6 trials. Shaded error bars are mean ± SEM. B) Reaching directions of cerebellar participants. C) Difference in reaching directions between cerebellar participants and control participants over all paradigms and phases of learning. Thin error bars indicate HDI of individual participants, thick error bars indicate HDI of group. % indicates percentage of HDI outside the ROPE. D) Difference in reaching directions between the standard paradigm and slow learning paradigms during the recovery phase of control participants. E) Difference in reaching directions between the standard paradigm and slow learning paradigms during the recovery phase of cerebellar participants.

Figure 4 shows that control participants learned the paradigms to approximately the same level. Learning was slower in the gradual paradigm because it was pinned to the gradual introduction of the perturbation in this paradigm. Learning also reached a very similar level in the different paradigms in the cerebellar participants, although it was slower and less complete than for control participants.

Behaviorally, the most salient difference between paradigms in control participants exists between the amount of spontaneous recovery in the standard paradigm versus the overlearning paradigm, i.e., there is more spontaneous recovery after overlearning than standard learning (Figure 4D). Our estimate of the difference between the overlearning paradigm and standard paradigm is 8.18° with a 95% HDI of l5.61°, 10.56°J, with more than 99.9% of the HDI outside the ROPE. In cerebellar participants, the difference between the overlearning paradigm and the standard paradigm also suggests more spontaneous recovery after overlearning (5.47° l3.05°, 8.06°J) though this difference is slightly smaller and obscured by the larger variability in the cerebellar patients (99.7% outside ROPE, Figure 4E). We compared the behavior of the subjects in the two groups. As expected, reaching directions of control participants and cerebellar participants are almost completely straight during the baseline phase (control group, all paradigms: −0.35° with 95% HDI of l-2.26°, 1.60°J, cerebellar group, all paradigms: −0.65° l-2.66°, 1.22°J). Movements in the baseline phase are quite similar between control and cerebellar participants (Figure 4C and supplementary statistical table).

When movements are perturbed by a visuomotor rotation, control participants learn the perturbation quickly (early adaptation, average across all paradigms excluding the gradual paradigm: 20.26° l17.72°, 22.19°J), and almost completely counteract the rotation late in the adaptation set (late adaptation, average across all paradigms: 26.08° l23.64°, 28.15°J). Cerebellar participants adapt more slowly and much less compared to control participants (early adaptation, average across all paradigms excluding the gradual paradigm: 12.63° l9.81°, 15.12°J, late adaptation, all paradigms: 18.17° l16.32°, 22.15°J). 99.9% of the HDI of the difference between control participants and cerebellar participants in early adaptation lies outside of the ROPE, suggesting that control participants really do learn faster; less than 99.97% of the HDI of the difference in late adaptation falls outside suggesting that control participants really do learn more fully (Figure 4C). This, of course, does not include the early adaptation phase of the gradual paradigm. Since the perturbation in the gradual paradigm in early adaptation averaged 4.2° and adaptation in the gradual paradigm was nearly linear, the final adaptation values in each group (26.05° and 18.21°) allow us to predict their early adaptation to be 3.6° and 2.5°. This gives a difference in early adaptation of only 1.1°. This is consistent with what is seen in the Figure 4C.

The spontaneous recovery phase produces similar values for control participants and cerebellar participants in all paradigms. The difference is near 0 but somewhat variable with 18%, 17%, 47%, and 56% outside ROPE for standard, gradual, overlearning and long ITI paradigms respectively (Figure 4C) meaning that we do not have strong evidence that spontaneous recovery is different between control and cerebellar participants in any of the paradigms.

We analyzed the possibility that our effects are driven by transfer of learning that depended on the order in which subjects did the different paradigms. The experiment was designed to address this possibility. First, after each learning phase, subjects did a wash-out block. In addition, subjects did the paradigms in 4 different orders, allowing us to assess the effect of the order of learning of the paradigms on the results. We did this by fitting two models of the order effects to the subject data and assessing both the strength of the order effects and their influence on our central results. This work is laid out in detail in the supplementary material. The results of this analysis show that our data is not sufficient to rule out the possibility of order effects on the order of 1-2°. However, we see robust increased spontaneous recovery in the overlearning paradigm in controls and, perhaps more weakly, in patients even when taking such order effects into account. In summary, out most salient behavioral result is that spontaneous recovery is increased in the overlearning paradigm in both cerebellar and control participants.

### 3.3 State-space Modelling

We reasoned that the additional spontaneous recovery in overlearning could have two different causes: increased slow learning at the end of adaptation or increased retention of slow learning during the counterperturbation. That is, prolonged activation of the slow system might increase the amount of slow learning. Alternatively, increased retention in the slow system could explain the results even with similar levels of slow learning. These hypotheses are explored in the next section.

Since behavioral differences between the paradigms were mainly evident in overlearning and standard learning, we focus on comparing these paradigms. The model results for the other paradigms can be found in the supplementary material.

Posterior predictive plots were used to assess the qualitative behavior of the model using the parameters estimated from the data (Figure 5). The model captures the structure of subject behavior: rapid learning is followed by a plateau and the short period of counterperturbation leads to spontaneous recovery. The model also captures the increased spontaneous recovery in the overlearning paradigm and the fact that this increase is somewhat smaller for cerebellar patients than for controls. The posterior predictive plots also support the hypothesis that spontaneous recovery reflects slow learning. For both groups, fast learning is essentially 0 during the spontaneous recovery phase (Figure 5A**, B, E,** and **F**).

**Figure 5:**
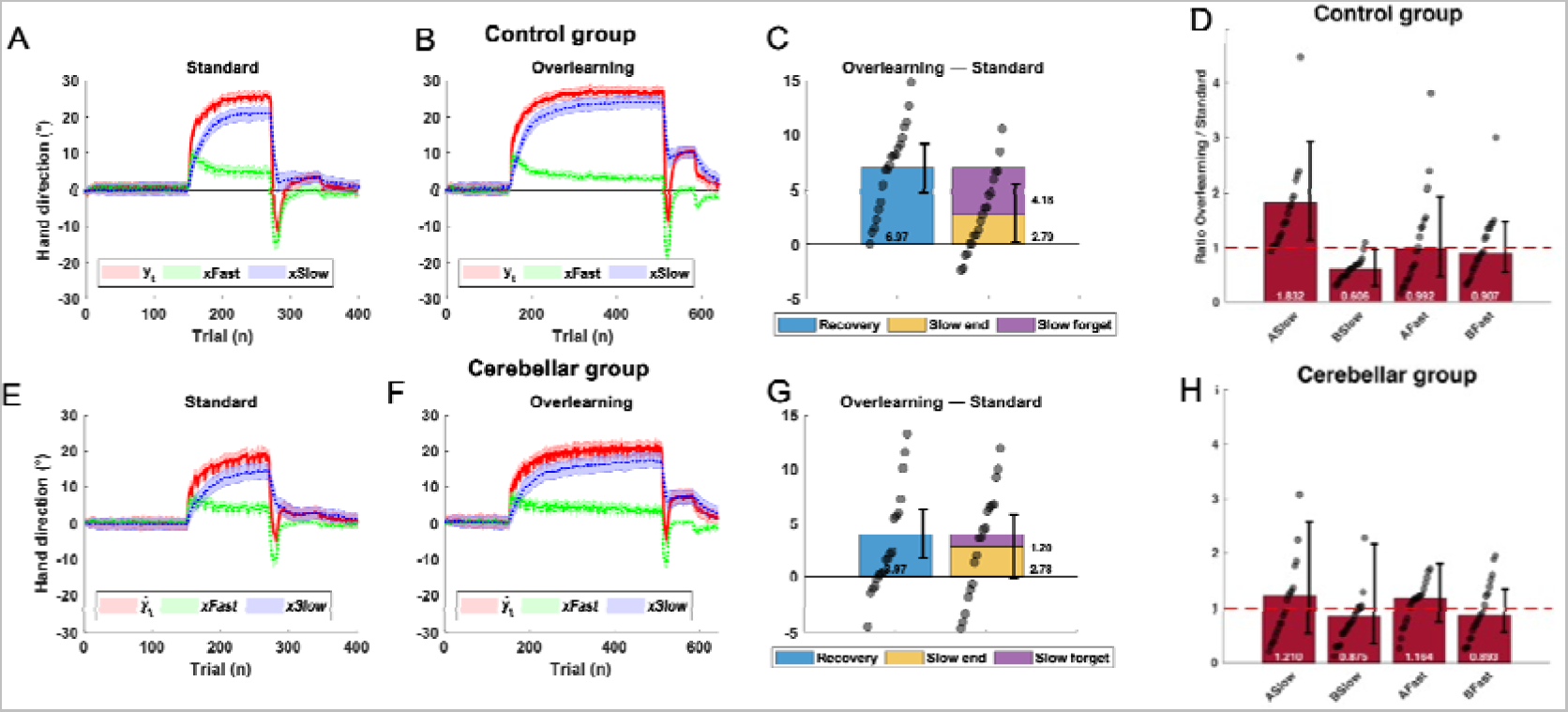
Posterior predictive plots of control participants in the A) standard and B) overlearning paradigm. The average model output ( ) is displayed with a solid red line, the average fast state ( ) with a dotted green line, and the average slow state ( ) with a dotted blue line. The shaded errorbars indicate the variability over posterior predictive samples around the average posterior predictive (95% HDI). C) shows the difference in spontaneous recovery of the slow system between the standard and overlearning paradigm (blue bar) and the difference in the amount of slow learning at the end of the adaptation set (yellow bar). The purple bar is the difference in the drop of slow learning after the counterperturbation trials between the standard and overlearning paradigm. Grey circles on the blue bar show results for individual subjects while the thick errorbar is the 95% HDI for the group. Grey circles on the yellow bar show results for individual subjects while the thick errorbar is the 95% HDI of the group. D) Change in learning and retention parameters between paradigms. The change in parameters is expressed as a ratio between the standard and overlearning paradigm. For A parameters the ratio was taken as ___________ , for B parameters the ratio was taken as ___________. This choice was made so that in both cases we are showing ratios of rates (forgetting rate and learning rate) and also so increased slow learning in overlearning would lead to larger numbers for both parameters. The gray circles represent ratios for individual participants, the thick black errorbar is the HDI of the population parameter. The red line indicates a ratio of 1. E), F), G), and H) display the same plots for cerebellar participants.

The spontaneous recovery exhibited by the model’s slow system in the two paradigms for the two groups is shown by the blue bars in Figure 5C **and 5G**. The difference between spontaneous recovery in the two paradigms in the model of the control participants is 6.97° l4.75°, 9.15°J (compared to 8.12° [5.67°, 10.66°] for actual participants); the difference for the model of cerebellar participants is 3.97° [1.65°, 6.22°] (compared to 5.60° [3.04°, 8.01°] for actual participants).

We hypothesized that spontaneous recovery is larger in overlearning due to either increased accumulation of slow learning or decreased forgetting in the slow system. Our modeling results suggest that both may be happening, but that they may be at different levels in the patients and control. This can be seen by comparing the yellow bar which shows increased accumulation of slow learning and the purple bar which shows decreased forgetting (increased retention) in the slow system (Figure 5C). Slow learning in controls is 2.79° [0.21°, 5.34°; 73% outside ROPE] higher in the overlearning paradigm than in the standard paradigm while slow forgetting (retention) is 4.18° [1.44°, 7.13°; 93% outside ROPE] more. Thus, for controls, about 60% of the increased spontaneous recovery comes from decreased forgetting and 40% from increased accumulation.

Difference in slow learning accumulation in controls is like that in cerebellar patients (2.77° [−0.22°, 5.71°; 70% outside ROPE]). In contrast, the difference in forgetting in patients is only 1.19° [−1.93°, 4.28°; 33% outside ROPE]. For cerebellar patients, 26% of the increased spontaneous recovery comes from decreased forgetting and 74% from increased accumulation.

Because of the apparent difference in how overlearning affects slow accumulation and forgetting in the two groups, we looked at the interaction. There was not much difference in the change in slow accumulation: 0.02° [−2.74°, 2.59°; 14% outside ROPE]. Controls had more reduced forgetting : 2.98° [−0.02°, 6.27°; 71% outside ROPE]. Taken together these results suggest that overlearning affected accumulation of the slow similarly in the two groups, and that forgetting changed in controls but probably not in patients. If there was a change in patients it was probably smaller than that in controls.

The rate of forgetting in controls in the counterperturbation is actually affected by four parameters of the model: the retention rate (A) and the learning rate (B) for both the slow and the fast states. Thus, decreased slow forgetting should be explained either by an increased retention rate (ASlow) or a decreased learning rate (BSlow). To explore this, we examined changes in learning and retention parameters between paradigms (Figure 5D**, H**). The figure suggests that in controls the slow retention parameter was larger in overlearning than in the standard paradigm (a ratio of 1.8 [1.1, 2.9]) and learning rate was smaller (ratio of 0.6 [0.3, 1.0]). This contrasts with retention and learning rate in the fast system, which are similar in the two paradigms (AFast: 1.0 [0.5, 1.9], BFast: 1.0 [0.6, 1.5]). In patients, all four parameters are similar in the two paradigms (ASlow: 1.2 [0.5, 2.6], BSlow: 0.9 [0.4, 2.2], AFast: 1.2 [0.8, 1.8], BFast: 0.9 [0.6, 1.3]).

The Supplementary Data includes an additional analysis where we used the model to show that additional training in the overlearning paradigm cannot explain our results if model parameters are held fixed.

#### Validating model results on participant data

The model suggests that overlearning causes a change in the parameters of the slow adaptation in control participants but not in cerebellar participants. This change in the parameters made slow adaptation slower to change, thus increasing spontaneous recovery. The analysis that follows is designed to validate the model.

We predicted that changes the model identified in slow learning and retention rates should affect the rate of adaptation. To test this, we measured the rate of adaptation in the counterperturbation phase, following adaptation, predicting that control subjects should have slower counterperturbation adaptation after overlearning than after the standard paradigm. This can be seen in Figure 6. Figure 6A shows that, for control participants, the slope of initial adaptation is not different between the overlearning and standard paradigms (overlearning: 1.73 [1.32, 2.09], standard: 1.73 [1.37, 2.14]). In contrast, Figure 6B shows that the slope of counterperturbation adaptation is reduced consistently across participants in the overlearning paradigm relative to the standard paradigm (overlearning: 2.71 [2.33, 3.11], standard: 3.13 [2.73, 3.50]). Figure 6C emphasizes this difference by plotting the difference between the slope in the standard and overlearning paradigms in the early adaptation and counterperturbation phases against each other (initial adaptation difference: 0.04 [−0.41, 0.5] with 67% outside ROPE; counterperturbation difference: −0.41 [−0.87, 0.05] with 92% outside of ROPE. Difference of differences: −0.48 [−1.10, 0.20] with 90% outside of ROPE). It is worth noting that a difference of 0.48 °/trial is a substantial difference in adaptation across tens of trials.

**Figure 6:**
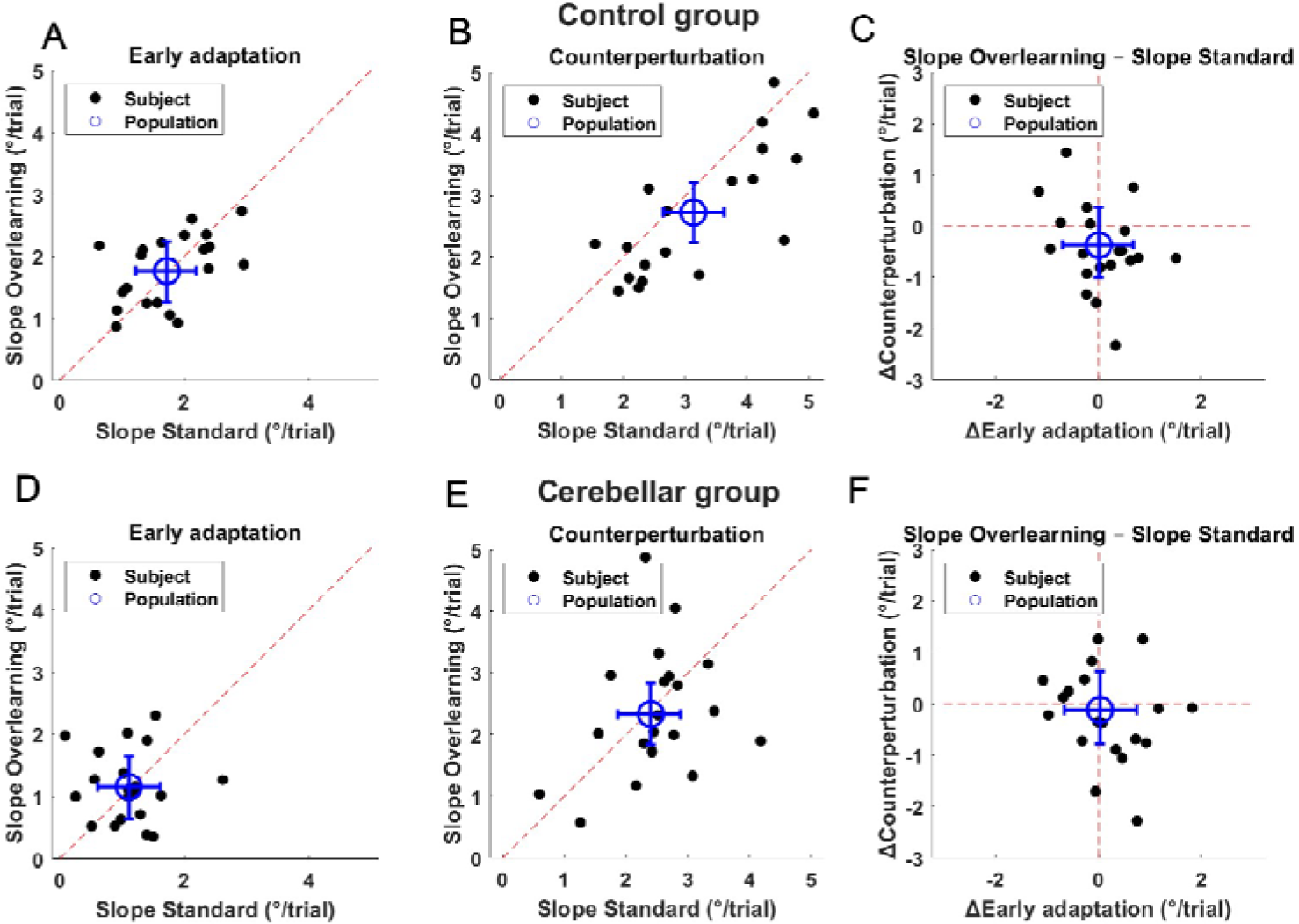
Comparison of the rate of adaptation in the early adaptation phase and the counterperturbation phase between the overlearning and standard paradigms. A) Slope of the adaptation in the first 13 trials of the adaptation in ovelearning and standard paradigms for control subjects. Dots show estimates for individual subjects. Blue cross shoes the estimate of the population. B) As in (A) but showing the slope of the adaptation in the first 13 trials of the counterperturbation phase. C) Difference in slope between overlearning and standard paradigms for the two different phases. D-F) Same format as A-C but for cerebellar participants.

Cerebellar patients show no similar effect. This can be seen by comparing Figure 6D to Figure 6B and Figure 6E to Figure 6C. Initial adaptation for patients was 1.07 [0.69, 1.49] in the standard paradigm and 1.13 [0.74, 1.55] in the overlearning paradigm (difference of 0.08 [−0.42, 0.54] with 70% outside of the ROPE). Counterperturbation adaptation was 2.43 [2.03, 2.83] in the standard paradigm and 2.30 [1.89, 2.70] in the overlearning (the difference was −0.12 [−0.62, 0.35] with 72% outside the ROPE). The difference of differences is −0.17 [−0.89, 0.49] with 80% outside of the ROPE.

## 4 Discussion

We tested 3 paradigms that have been thought to increase slow learning in both controls and cerebellar patients. For each we examined spontaneous recovery after a short counterperturbation, a phenomenon that has been taken as a hallmark of slow learning (Smith et al., 2006). We find that only one paradigm, overlearning, increased spontaneous recovery and that it did so in both controls and cerebellar patients (Figure 4). Intuitively, increased spontaneous recovery might been taken as a sign that the level of slow learning is higher at the end of adaptation. We tested this by fitting a two-state model of motor adaptation to our data using a Bayesian fitting procedure and generating posterior predictive data to model the behavior of the subjects. Our model showed increased spontaneous recovery, especially in control subjects (Figure 5). However, this did not result from a higher level of slow learning at the end of adaptation. Rather, overlearning led to a smaller drop in the slow learning during the counterperturbation. This seemed to be because, in controls, the model parameter for retention of slow learning (ASlow) was higher during overlearning than during the standard paradigm and the model parameter for slow learning rate (BSlow) was lower during the overlearning than in the standard paradigm. To validate these results, we tested a post-hoc prediction that these changes in the parameters of learning would slow adaptation. We tested this by comparing the adaptation rate during the counterperturbation in the standard paradigm and the overlearning paradigm. The adaptation was, indeed, slower in controls, a result that was not reproduced in the cerebellar patients.

Our primary result is that the overlearning task was the only paradigm that led to more spontaneous recovery than the standard tasks. There was logic to thinking the other tasks would also lead to greater spontaneous recovery. Gradual adaptation has long been thought to decrease the rate at which retention washes away compared to abrupt adaptation (Criscimagna-Hemminger et al., 2010; Huang and Shadmehr, 2009; Kagerer et al., 1997; Kluzik et al., 2008; Michel et al., 2007; Wong and Shelhamer, 2011) although there have been some studies that did not find such an effect (Klassen et al., 2005; Werner et al., 2014). A more recent paper suggests that the apparent effects on retention in the gradual condition actually reflects secondary effects either of training duration or level of learning (Alhussein et al., 2019). This is entirely consistent with our results. Our expectation regarding an effect of long ITIs is rooted in the history showing increased retention following spaced learning compared to massed learning (for a review see Smolen et al., 2016), that has been documented long ago also in motor adaptation (Taub and Goldberg, 1973). Indeed, recent work has suggested that long ITIs should drive stronger retention in adaptation (Kim et al., 2015; Zhou et al., 2017). However, in these studies, effects are small and individual subject variability is large with delays on the order of 15 sec. In both studies, details of the experimental protocol were also quite different from our own. Thus, it might be possible to see effects with a long ITI, but it would require delays that are quite long and a different task. In sum, our results are consistent with other results in the literature raising doubts about whether either gradual adaptation or long ITIs drive the slow learning system.

Overlearning – the continued practice once performance has plateaued – has long been known to increase retention (for review see Driskell et al., 1992). The effect of the number of training trials on retention of motor adaptation has been confirmed more recently (Joiner and Smith, 2008; Yamada et al., 2019). Indeed, as mentioned, Alhussein (2019) showed that previously reported effects of gradual adaptation on slow learning are, to a large extent, mediated by the number of training trials. Our findings are consistent with these previous findings. However, the effects shown previously are different from ours. First, Joiner and Smith (2008) and Yamada (2019) both show increased retention after 24 hours, while Alhussein (2019) measures the rate of decay of performance in a no feedback condition immediately after training. Ours is the first paper to our knowledge to show an effect of overlearning on spontaneous recovery.

Our second basic result is that spontaneous recovery was also increased in patients, although in a manner that was less pronounced than in controls. Consistent with many previous findings, cerebellar subjects learned slower than controls and reached a lower level of adaptation at the end of the adaptation (Figure 4). During spontaneous recovery, their performance was close to that of controls for all paradigms, including the overlearning. This suggests that overlearning does cause some increased slow learning in patients as well as controls.

Our third basic result comes from our modeling which suggests that overlearning changes the parameters of the slow system in controls. Joiner and Smith (2008) ascribe most of the effect of overlearning to an increase in the level of slow adaptation with increased number of training trials. This contrasts with our interpretation: most of the effect reflects slow adaptation becoming more resistant to change as overlearning progresses. The differences may arise from the length of training in the two studies. Joiner and Smith (2008) show an effect that begins to level off near 100 training trials and the group with the most training only does 160 training trials. Our study had one group with around 108 training trials and another with 324 training trials. Thus, it is possible that what they see reflects a slow rise in the level of the slow adaptation system, but that after this system reaches a plateau increased training begins to change its responsiveness.

Model results also suggest that at the end of the standard paradigm, the slow system of cerebellar patients has not yet reached plateau and some increase is still possible in the slow learning system. The model suggests that cerebellar patients have less slow learning than controls and more slow forgetting. Fast learning of cerebellar patients and controls is quite similar, consistent with earlier experimental findings (Taylor et al., 2010; Wong et al., 2019). The idea that the history of adaptation influences its dynamics is not new (Shadmehr and Brashers-Krug, 1997). Our suggestion that overlearning may lead to increased resiliency in the slow system is in line with previous research specifically suggesting that environmental consistency affects adaptation dynamics (Avraham et al., 2019). Indeed, previous work has also used changing fit of the two state model as evidence of changes in learning parameters (Mawase et al., 2014). The validation of our results provided by the post-hoc analysis further underlines these points.

Conclusions drawn from our research must be tentative due to a number of limitations. One important caveat is that our design is a within subjects design with each subject doing all tasks on the same day. We addressed this limitation in different ways. First, we used washout at the end of each session to reduce transfer (Caithness et al., 2004; Krakauer et al., 2005; Nguyen et al., 2019; but see also Kitago et al., 2013; Villalta et al., 2015). Second, order of the tasks and perturbation direction was counterbalanced between subjects and we tested for order effects (see supplementary materials). We do not believe that our central result is sensitive to possible inter-session effects. A second concern is that subtle details of the task – such as the fact that the gradual adaptation group did not spend any time in the plateau or that the long ITIs may have been too short -- may have specifically influenced our results. This possibility cannot be ruled out. A third concern is one that is characteristic of all patient studies: patients are variable, disease etiology is complex, and we are specifically studying chronic lesions.

Perhaps the most salient limitation of our study is in the interpretation of the modeling. The modeling work cannot be conclusive, and alternative explanations exist. Overlearning may engage additional learning mechanisms that are not accounted for by the two-state model. For instance, Therrien et al., (2016) has shown that reinforcement learning can be selectively engaged in the adaptation task, and that this can affect the way subjects behave during error clamps (Shmuelof et al., 2012). Additional mechanisms to consider are use-dependent learning (Diedrichsen et al., 2010) or model-free learning (Huang et al., 2011). Finally, a recently published model uses dynamic formation of specific adaptation memory to explain many of the dynamics of adaptation (Heald et al., 2020). Thus, our findings and modeling work lend credence to the idea that extra practice makes learning more resilient, but they do not conclusively identify the mechanisms based on comparison of alternative models.

In sum, our research shows that spontaneous recovery is specifically affected by overtraining in cerebellar patients and controls. We hypothesize that, in controls, this is primarily driven by changes in the dynamics of slow adaptation while in cerebellar patients it reflects changes in the amount of slow adaptation achieved. That is, while residual slow learning does exist in cerebellar patients, we do not see evidence that the primary cerebellar deficit can be improved by overtraining paradigms.

## Acknowledgements

The study was funded by grants of the German Research Foundation (DFG TI 239/16-1 awarded to OD and DT; DFG project number 316803389 – SFB 1280 [subproject A05] awarded to DT), and a scholarship from the Essener Ausbildungsprogramm “Labor und Wissenschaft” für den Aerztlichen Nachwuchs (ELAN) supported by the Else Kröner-Fresenius-Stiftung awarded to AM. We would like to thank Beate Brol for her support in the analysis of this experiment. We are deeply grateful to Jonathan Tsay for his thoughtful discussion and comments.

