## Supplementary materials for "Can we improve slow learning in cerebellar patients?"

**Supplementary materials: Can we improve slow learning in cerebellar patients?**

**State space modeling**

A two-state model was fit to the reaching directions of all trials in each individual participant. The equations for the state-space model are given in Equations 1$-6$:

$e_{t}=y_{t}+p_{t}$	(Eq. 1)

$x_{t+1}^{F}=A^{F}x_{t}^{F}-B^{F}e_{t}+\epsilon_{state}$	(Eq. 2)

$x_{t+1}^{S}=A^{S}x_{t}^{S}-B^{S}e_{t}+\epsilon_{state}$	(Eq. 3)

$x_{t}=x_{t}^{F}+x_{t}^{S}$	(Eq. 4)

$y_{t}=x_{t}+\epsilon_{output}$	(Eq. 5)

$\epsilon_{output} \sim N\left( 0,\sigma_{\epsilon_{o}}^{2} \right), \epsilon_{state}\sim N(0,\sigma_{\epsilon_{s}}^{2})$	(Eq. 6)

Where $A^{S}>A^{F}$ and $B^{S}< B^{F}$.

This state-space model, as posited by Smith and colleagues (Smith et al., 2006), consists of a fast state ($x_{t}^{F}$) and slow state ($x_{t}^{S}$). The fast state learns quickly and forgets quickly, while the slow state learns slowly and forgets slowly. Both states have an independent learning rate ($B$) and retention rate ($A$). After each movement, the fast and slow state are updated based on the error ($e)$ in the previous movement. In visuomotor experiments, the amount of error is given by addition of the actual movement ($y_{t}$) and perturbation ($p_{t}$).

To estimate the parameters and hidden state variables for a given participant from reaching directions, we implemented a hierarchical model based on Equations $1-6$ in JAGS (**Supplementary** **Figure 1**). A hierarchical model improves fits by sharing information across participants. The parameters and state variables were estimated by MCMC methods, separately for each of the four training paradigms (standard, gradual, overlearning and long ITI) and two participant groups (cerebellar participants or control participants). In the hierarchical model, a participants’ learning rate ($B^{S}$ and $B^{F}$) and retention rate ($A^{S}$ and $A^{F}$) came from a normal distribution centered around $\mu$ with precision $\tau$. The learning and retention rates were sampled in logistic space and then transformed to the range from 0$-1$ to more realistically reflect changes in the parameters and provide better sampling behavior. Precision for the states and execution noise came from a very broad gamma distribution ($\text{A}={10}^{-3}, \text{B}= {10}^{-3}$).

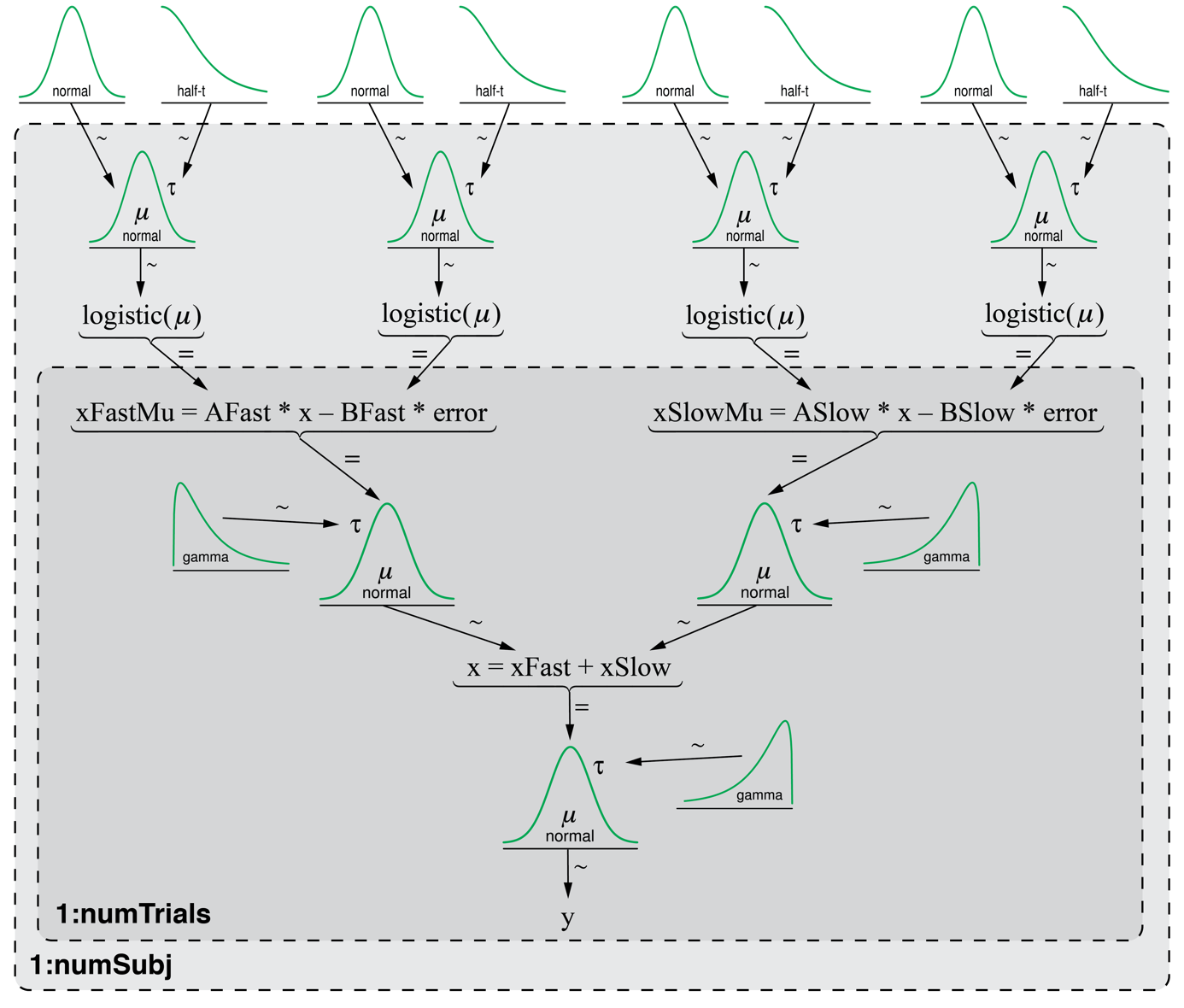

**Supplementary Figure 1**: Diagram of the hierarchical model implemented in JAGS. The equals sign (=) denotes deterministic relationships, while tildes (~) denote stochastic relationships. The distributions on the white background indicate the hyperpriors. The light grey background indicates the subject loop, while the dark grey background indicates the trial loop.

Only the shape of the learning- and retention rate distributions was specified (normal in logistic space), the actual priors for each participant were sampled from a hyperprior. The hyperparameter for $\mu$ came from an uninformative normal distribution ($M=0, S=1000$) and $\tau$ from a weakly informed half-t distribution ($M^{'}=0, S^{'}=2.5, N=7$) so each posterior distribution was mainly informed by the data. Reasonable initial values for $M$ and $S$ were estimated by running the model first without hyperpriors and weakly informed priors for the learning and retention rates of a participant (4 chains, adaptation and burn-in of 10,000 samples, 30,000 samples collected). Model runs for the model with hyperparameters started with an adaptation phase of 20,000 samples and burn-in phase of 80,000 samples, after which we collected 250,000 samples on four separate chains. Samples were thinned by a factor of 5 to decrease the memory footprint and autocorrelation of samples. For every parameter we first visually inspected the trace plots for chain convergence. Next, several MCMC diagnostics were calculated: the ESS, PSRF and MCSE (**Supplementary Table 1**). Generally, sampling of the posterior distributions was considered adequate. No large differences in ESS, PSRF and MCSE were found between the paradigms. A small number of parameters had low ESS due to high autocorrelations in specific paradigms, mainly on the participant level, but longer model runs were deemed impractical. **Supplementary table 2** displays the estimated modal values and HDI’s of the hyperparameters of both control participants and cerebellar participants in all paradigms.

| **Supplementary table 1** | | | | | | |
| --- | --- | --- | --- | --- | --- | --- |
| *MCMC diagnostics* | | | | | | |
| **Cerebellar participants** | | | | **Control participants** | | |
| **Parameters** | **ESS** (Q1 – Q3) | **PSRF** | **MCSE** | **ESS** (Q1 – Q3) | **PSRF** | **MCSE** |
| ASlowHypMu [4] | 4,296 (3,213 – 5,297) | 1.0010 | <0.0001 | 2,789 (1,178 – 4,972) | 1.0098 | 0.0003 |
| AFastHypMu [4] | 1,149 (829 – 1,487) | 1.0017 | 0.0002 | 957 (422 – 1,685) | 1.0053 | 0.0006 |
| BFastHypMu [4] | 8,581 (6,536 – 12,557) | 1.0004 | <0.0001 | 39,094 (11,205 – 74,919) | 1.0001 | <0.0001 |
| BSlowHypMu [4] | 2,231 (1,692 – 2,558) | 1.0039 | 0.0004 | 1,181 (561 – 1,487) | 1.0051 | 0.0004 |
| ASlowHypPrec [4] | 3,490 (2,970 – 4,440) | 1.0013 | 0.0002 | 3,697 (1,459 – 41,582) | 1.0023 | 0.0005 |
| AFastHypPrec [4] | 1,371 (9,49 – 1,484) | 1.0008 | 0.0010 | 1,266 (709 – 1,764) | 1.0028 | 0.0010 |
| BFastHypPrec [4] | 4,750 (2,649 – 7,278) | 1.0007 | 0.0003 | 19,099 (7,516 – 37,255) | 1.0005 | <0.0001 |
| BSlowHypPrec [4] | 3,911 (2,150 – 5,198) | 1.0015 | 0.0004 | 2,332 (1,102 – 2,954) | 1.0049 | 0.0023 |
| ASlow [4,20] | 7,684 (2,990 – 14,159) | 1.0011 | <0.0001 | 6,119 (2,340 – 12,758) | 1.0024 | <0.0001 |
| AFast [4,20] | 1,430 (894 – 2,259) | 1.0024 | 0.0001 | 803 (386 – 1,249) | 1.0055 | 0.0003 |
| BSlow [4,20] | 3,317 (1,365 – 8,521) | 1.0018 | <0.0001 | 1,001 (484 – 2,089) | 1.0059 | <0.0001 |
| BFast [4,20] | 2,493 (1,572 – 4,009) | 1.0012 | <0.0001 | 2,217 (943 – 4,339) | 1.0021 | <0.0001 |
| outputPrec [4,20] | 12,867 (6,543 – 24,204) | 1.0004 | 0.0001 | 33,614 (19,436 – 50,775) | 1.0003 | <0.0001 |
| statePrec [4,20] | 1,857 (970 – 2,998) | 1.0019 | 0.0906 | 2,835 (1,420 – 5,568) | 1.0019 | 0.0818 |

**Supplementary Table 1**: Parameter names correspond with nodes in JAGS model code. For brevity, the median ESS, mean PSRF and mean MCSE taken over the four different paradigms are displayed. Numbers in brackets indicate the levels per parameter. Note that the model was run separately for each of the paradigms, i.e. the first level relates to the four chains of one paradigm. The interquartile range (Q1 – Q3) of the ESS is given between parentheses.

| **Supplementary table 2** | | | | | | | | |
| --- | --- | --- | --- | --- | --- | --- | --- | --- |
| *Mode and HDI’s of hyperparameters in two-state model* | | | | | | | | |
| **Control participants** | | | | | | | | |
|  | **Standard** | | **Gradual** | | **Overlearning** | | **Long ITI** | |
| **Parameter** | **Mode** | **HDI** | **Mode** | **HDI** | **Mode** | **HDI** | **Mode** | **HDI** |
| ASlowHypMu | 0.992 | 0.989 – 0.994 | 0.998 | 0.994 – 1.000 | 0.996 | 0.994 – 0.997 | 0.992 | 0.988 – 0.995 |
| AFastHypMu | 0.811 | 0.664 – 0.903 | 0.850 | 0.783 – 0.894 | 0.820 | 0.750 – 0.868 | 0.847 | 0.725 – 0.910 |
| BFastHypMu | 0.162 | 0.113 – 0.221 | 0.187 | 0.116 – 0.277 | 0.153 | 0.106 – 0.214 | 0.138 | 0.095 – 0.198 |
| BSlowHypMu | 0.060 | 0.040 – 0.103 | 0.043 | 0.028 – 0.068 | 0.038 | 0.029 – 0.052 | 0.046 | 0.033 – 0.068 |
| **Cerebellar participants** | | | | | | | | |
|  | **Standard** | | **Gradual** | | **Overlearning** | | **Long ITI** | |
| **Parameters** | **Mode** | **HDI** | **Mode** | **HDI** | **Mode** | **HDI** | **Mode** | **HDI** |
| ASlowHypMu | 0.989 | 0.982 – 0.994 | 0.992 | 0.982 – 0.998 | 0.992 | 0.986 – 0.995 | 0.990 | 0.982 – 0.995 |
| AFastHypMu | 0.587 | 0.454 – 0.696 | 0.647 | 0.532 – 0.735 | 0.667 | 0.550 – 0.757 | 0.631 | 0.527 – 0.716 |
| BFastHypMu | 0.149 | 0.115 – 0.188 | 0.174 | 0.130 – 0.230 | 0.134 | 0.094 – 0.185 | 0.154 | 0.118 – 0.194 |
| BSlowHypMu | 0.019 | 0.013 – 0.029 | 0.018 | 0.009 – 0.030 | 0.019 | 0.009 – 0.040 | 0.009 | 0.005 – 0.016 |

**Supplementary Table 2**: Mode and HDI of credible parameter values in control participants and cerebellar participants. HDI is the 95% highest density interval. Parameter names correspond with nodes in JAGS model code.

**Model results: switched parameters and paradigms**

The change in the model behavior in the overlearning task is likely caused by the changed parameters as discussed in the main text. However, it is also possible that changes in model behavior reflect the extended adaptation phase in the overlearning paradigm. To test the effect of the changes in the model parameters on model behavior, we generated posterior predictive data for the overlearning paradigm using parameters from the standard paradigm and for the standard paradigm using parameters from the overlearning paradigm. The results of this are shown in **Supplementary** **Figure 2**. The figure shows that control participants have smaller spontaneous recovery in the overlearning paradigm when using parameters from the standard paradigm (**Supplementary Figure 2A**). The change in spontaneous recovery caused by this parameter switch is -7.4° [-10.9°, -4.2°] (comparing spontaneous recovery in **Supplementary Figure 2A** and **Main Text Figure 5B**). Similarly, spontaneous recovery in the standard paradigm is larger using parameters from the overlearning paradigm (3.8° [0.8°, 7.0°]) (comparing **Supplementary Figure 2B** and **Main Text Figure 5A**). Thus, in controls, the change in the parameters caused by the overlearning seem to explain the difference in spontaneous recovery better than just the overlearning itself.

We further ask whether the changed parameters cause increased levels of slow learning during adaptation or a reduced forgetting during the counterperturbation. For control participants, the final level of slow learning at the end of adaptation is not very different when the parameters are changed (certainly for the difference in the standard paradigm, **Supplementary Figure 2A** and **Main Text Figure 5B**: 0.5° [-2.9°, 3.5°], but also in the overlearning paradigm, **Supplementary Figure 2B** and **Main Text Figure 5A**: 2.1° [-0.6°, 4.9°]). In contrast, the effects of parameters on slow forgetting in the counterperturbation is more pronounced (in the standard paradigm: 3.3° [-0.6°, 7.4°] and in the overlearning: 5.4° [4.7°, 9.3°]).

The effects of the parameters on the final level of slow adaptation and on slow forgetting are most clearly demonstrated in **Supplementary Figure 2C** which shows the differences by comparing the two paradigms when the parameters are switched (**Supplementary Figure 2A** and **7B**)**.** Switching parameters between paradigms leads to a change in spontaneous recovery (4.2° [0.4°, 8.2°]) that is almost completely explained by slow forgetting (4.5° [0.3°, 9.1°]) and does not depend at all on slow learning (-0.3° [-3.6°, 3.3°]). However, while these differences are striking, they are below nominal levels of significance (percentage of the posterior distribution outside the ROPE: spontaneous recovery: 87%, slow learning: 27%, slow forgetting: 86%).

We did a similar analysis of the model fit to cerebellar patients (**Supplementary Figure 2D-F**). Here, the difference of using standard parameters in the overlearning paradigm also caused a decrease in spontaneous recovery, although smaller than that in control (**Supplementary Figure 2D** and **Main Text Figure 5E**, -2.6° [-6.1°, 1.0°]). Similarly, switching parameters in the standard paradigm slightly increased spontaneous recovery (**Supplementary Figure 2E** and **Main Text Figure 5D**, 1.6° [-1.4°, 4.8°]). These shifts in spontaneous recovery lead to nearly identical spontaneous recovery across paradigms when the parameters are switched (**Supplementary Figure 2C**; 0.3° [-3.9°, 4.3°]). In line with this, slow adaptation and slow forgetting seem quite similar when the parameters are swapped across paradigms (slow learning: 0.6° [-3.4°, 4.6°]; slow forgetting -0.9° [-5.4°, 4.0°]).

To sum up our model results, increased spontaneous recovery in controls is explained by the model as a change in the parameters of the slow system. No evidence for such a change in parameters is seen in the patients.

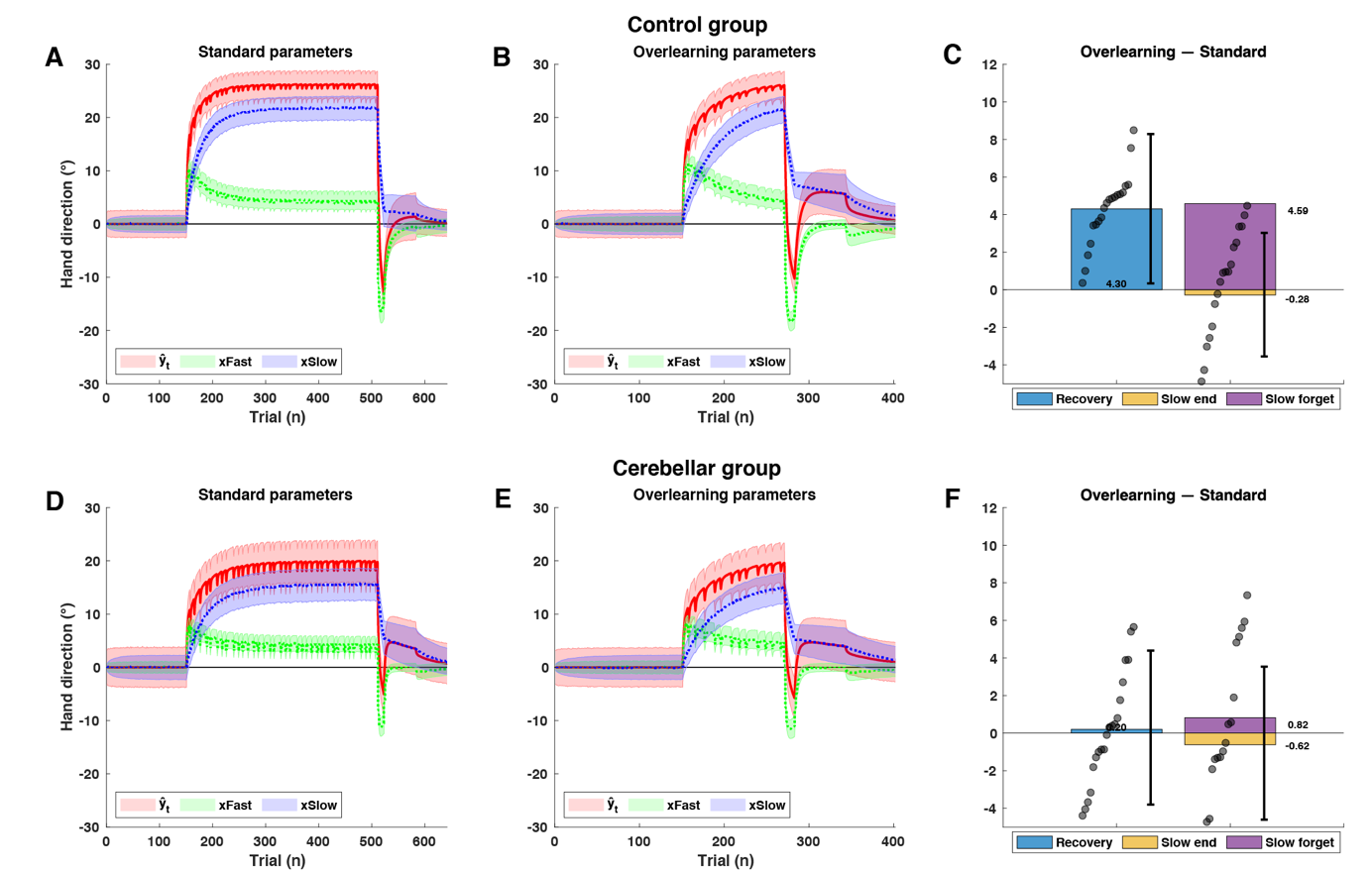

**Supplementary Figure 2:** Posterior predictive plots generated using incongruent parameters. That is, the posterior predictive plots for the overlearning paradigm were generated from the parameters fit to the standard paradigm and vice versa. A) Control participants with standard parameters in the structure of the overlearning paradigm and B) overlearning parameters in the structure of the standard paradigm. The average model output ($\hat{y}_{t}$) is displayed with a solid red line, the average fast state ($x_{t}^{F}$) with a dotted green line, and the average slow state ($x_{t}^{S}$) with a dotted blue line. The shaded errorbars indicate the variability around the average posterior predictive (2.5^th^ percentile – 97.5^th^ percentile of simulated data). Panel C) shows the difference in spontaneous recovery of the slow system between the standard parameters and overlearning parameters (blue bar) and the difference in the amount of slow learning at the end of the adaptation set (yellow bar). The purple bar is the difference in the drop of slow learning after the counterperturbation trials between the standard and overlearning parameters. Grey circles on the blue bar indicate individual differences in spontaneous recovery, the thick errorbar is the HDI of the group. Grey circles on the yellow bar indicate individual differences in slow learning at the end of adaptation, the thick errorbar is the HDI of the group. Panels D), E) and F) display the same plots for cerebellar participants instead.

**Model results: gradual and long ITI paradigms**

Posterior predictive plots were generated for the gradual and long ITI paradigm as well. **Supplementary Figures 3 and 4** are in the format of **Main Text Figure 5**, but the gradual (**Supplementary Figure 3)** and long ITI paradigm (**Supplementary Figure 4**) are depicted instead. The predicted difference in spontaneous recovery between the gradual and standard paradigm, as well as the long ITI and standard paradigm, is smaller than the predicted difference between the overlearning and standard paradigm for both groups (cf. **Main Text Figure 5**). This is probably best understood as a result of the reduced slow learning at the end of adaptation in both groups. Both groups reach a balance of slow learning and fast learning at the end of the counter-perturbation that is very similar to the one in the standard group. Examination of the ratio of parameters between groups shows that the most striking difference between the overlearning and the other groups is the increase in ASlow in the control participants (**Supplementary Figure 5 and 6 and Main Text Figure 5**).

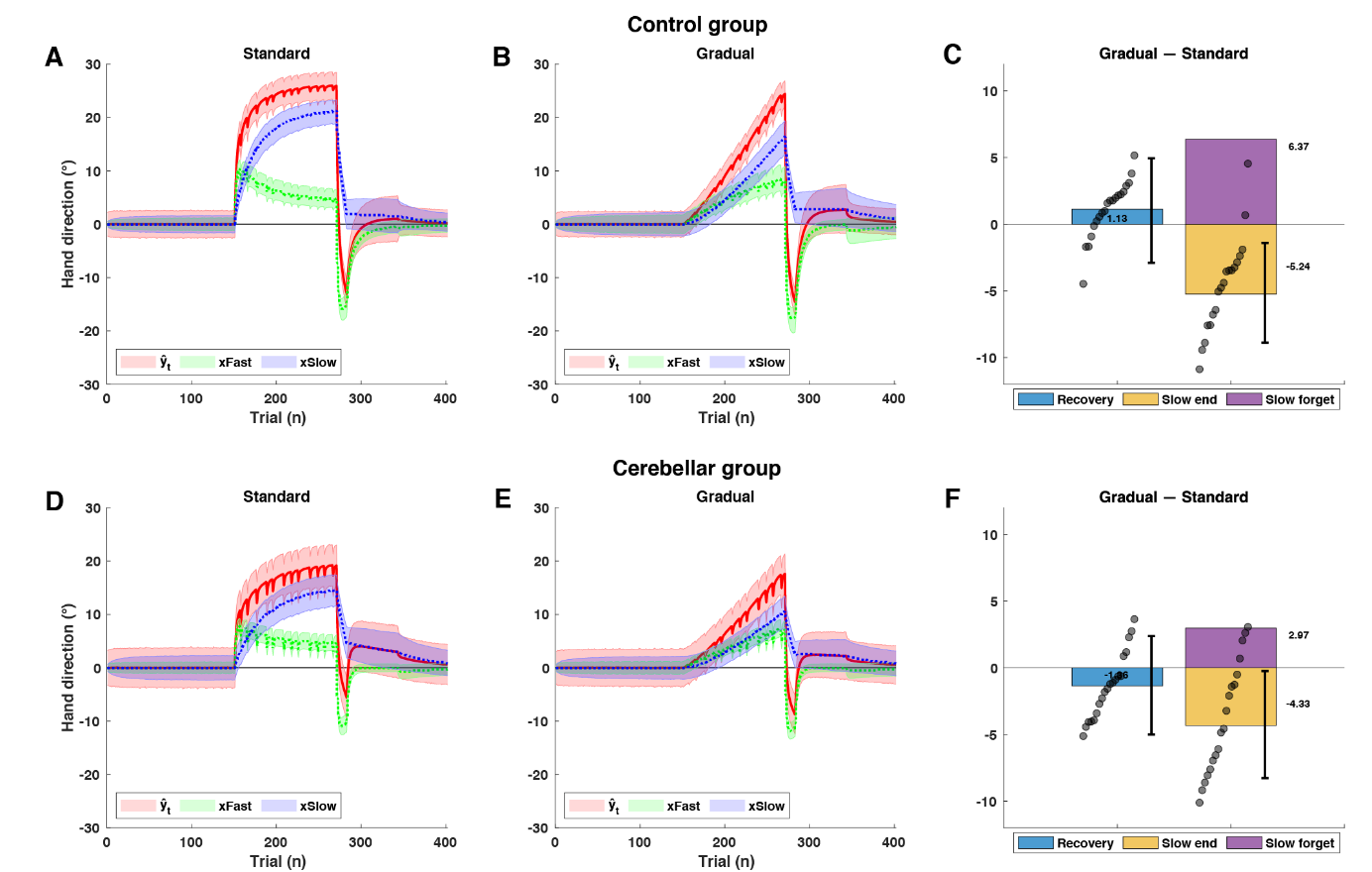

**Supplementary figure 2**: Posterior predictive plots comparing the standard and gradual paradigm. Cf. **Main Text Figure 5**.

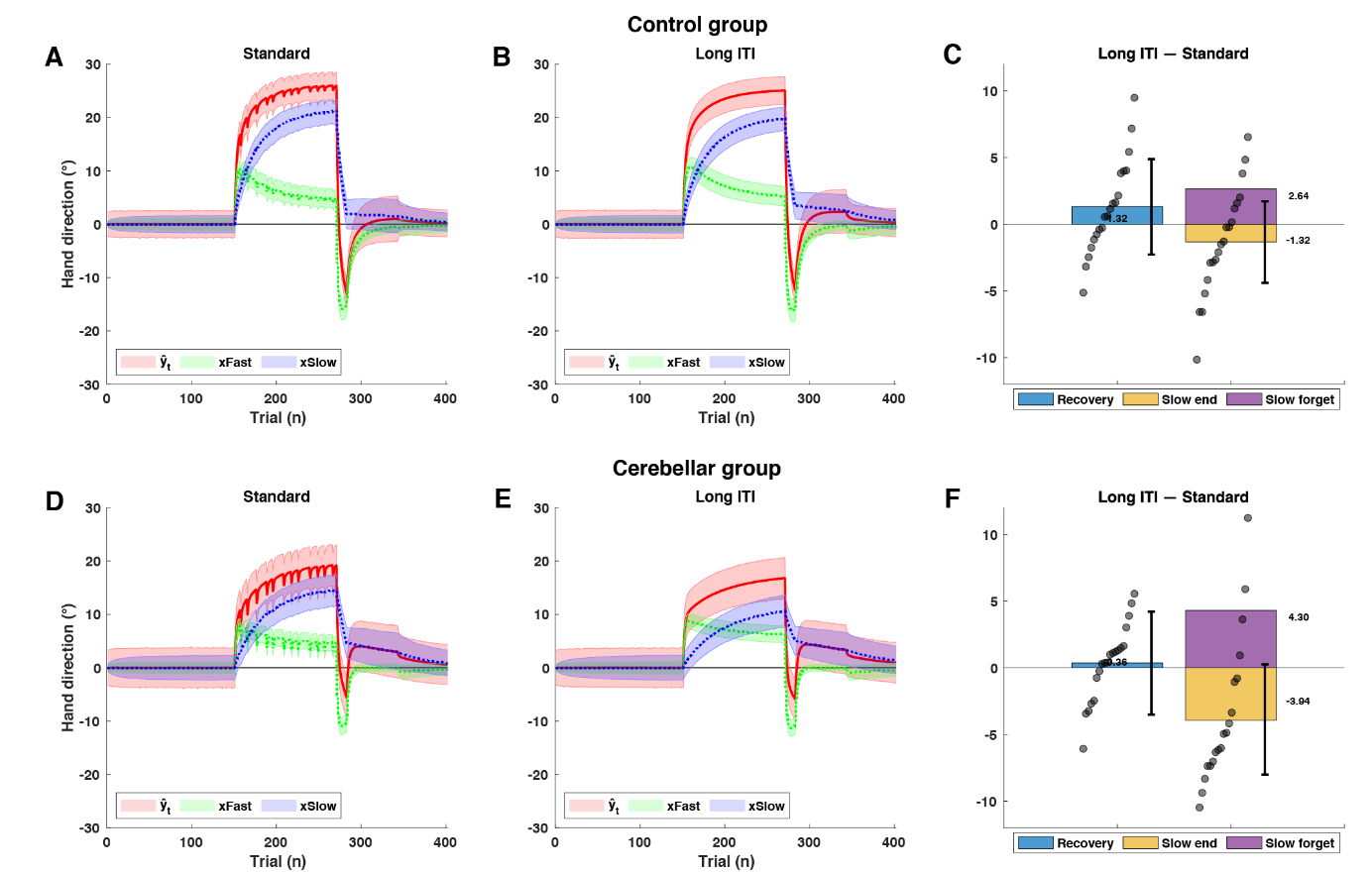

**Supplementary Figure 4**: Posterior predictive plots comparing the standard and long ITI paradigm. Cf. **Main Text Figure 5** of the main text.

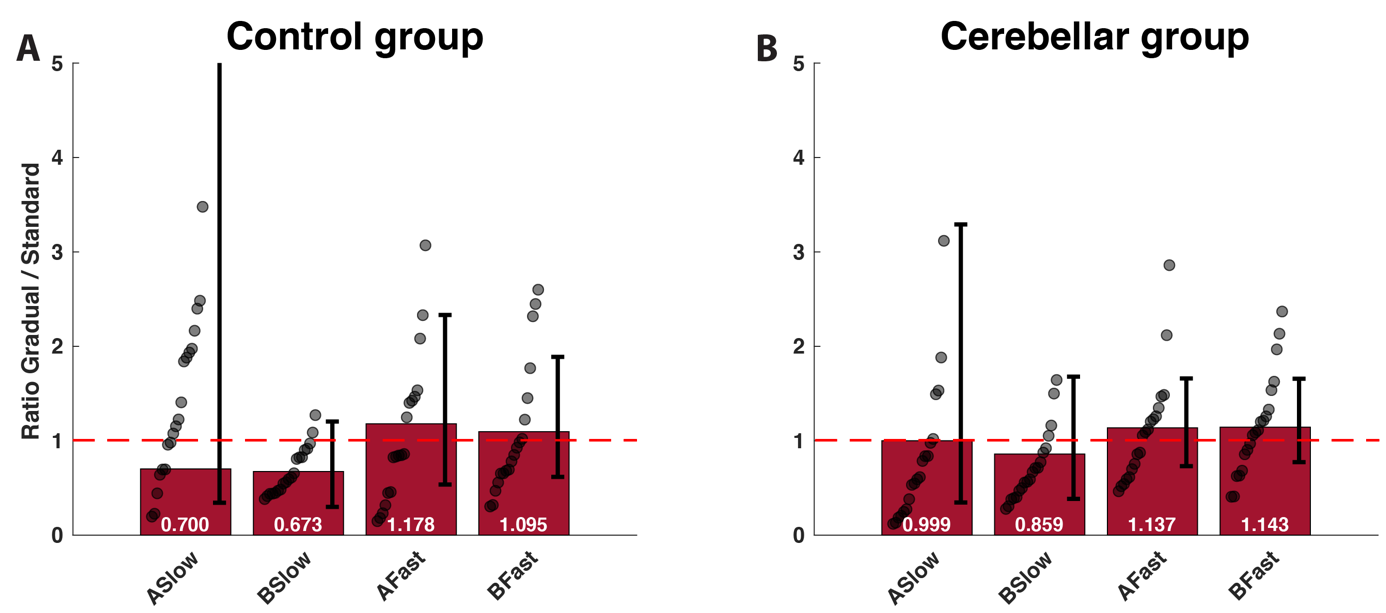

**Supplementary Figure 5**: Change in learning and retention parameters between gradual and standard paradigm. Cf. **Main Text Figure 5**.

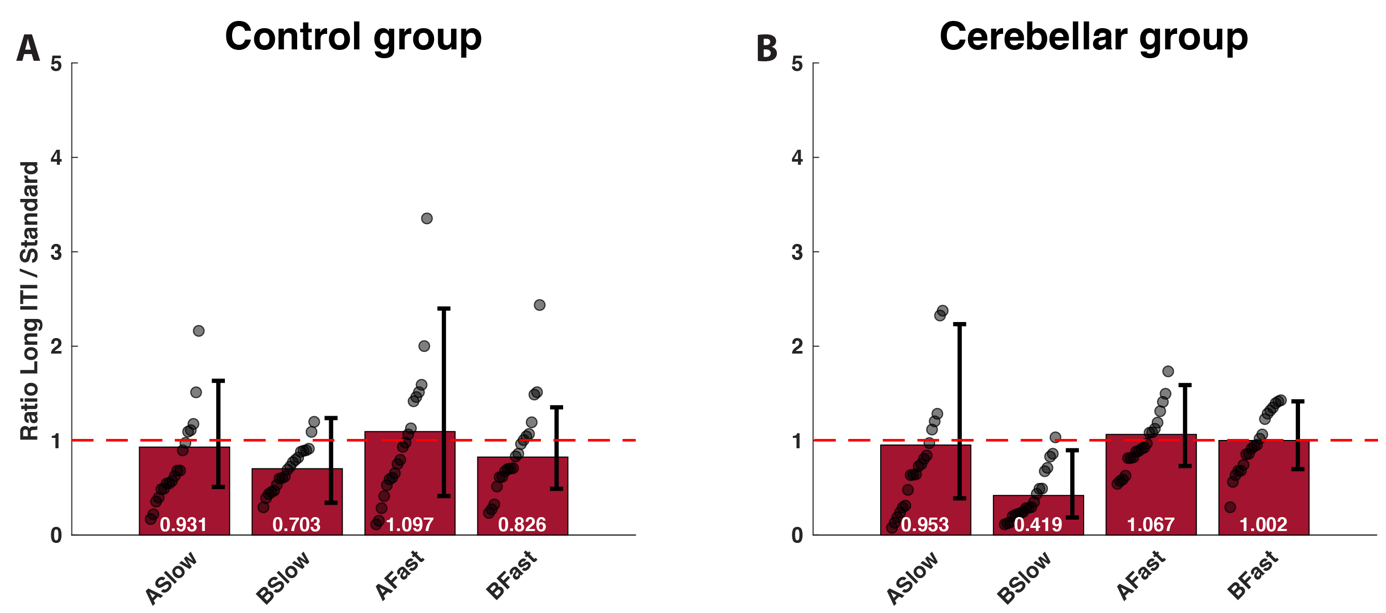

**Supplementary Figure 6**: Change in learning and retention parameters between long ITI and standard paradigm. Cf. **Main Text Figure 5**.

**Order effects**

Subjects did the 4 paradigms in 4 different orders chosen using a latin squares design such that there were 5 subjects in each group doing each particular order of paradigms. To assess order effects, we redid that regression analysis described in the main text but added an additional factor for order effects. That is, the results for each subject were specified as:

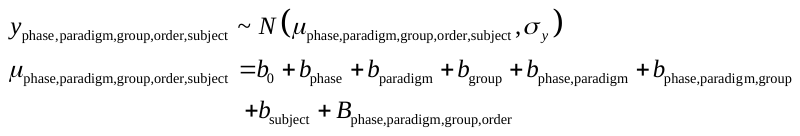

Where the
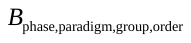
 term was determined by the specific paradigms that preceded the current paradigm in the current order. We did this following two different models for order effects (**Supplementary Figure 7**). In the first model (**Supplementary Figure 7A**), order effects were persistent: the effect of doing one paradigm on all subsequent paradigms was the same. In the second model (**Supplementary Figure 7B**), only the immediately preceding paradigm affected performance on the current paradigm. That is, we included regression coefficients for each phase group and preceding paradigm and calculated the term
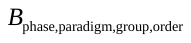
 as either

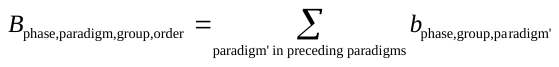

or

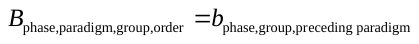

The full model is available at: https://github.com/Motor-Learning-Lab/paper-slowlearning

The results of this analysis show that all estimated order effects in both models were distributed relatively symmetrically around 0 with means of 0.17 (immediate model) and 0.09 (persistent models) and standard deviations of 1.5° (immediate) and 1.7° (persistent). None of them had less than a 40% overlap with the ROPE and 38 out of 64 estimates of the order effects across both models (59%) had an overlap of greater than 95% with the ROPE indicating values that are practically equivalent to 0. Relaxing the constraint, 52 out of 64 (81%) had an overlap of greater than 80% with the ROPE. Thus, overall, while we cannot rule out the presence of order effects given our data, we also did not see any evidence to support their existence. Focusing specifically on the order effects during the spontaneous recovery period, the maximum estimated order effect was for an increase of up to 3.8° [ROPE:
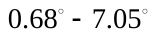
, 41% overlap with ROPE] degrees in spontaneous recovery for paradigms that followed the overlearning paradigm in control patients (using the persistent model for order effects). This particular order effect, if it is real, would tend to reduce our estimate of the effect and could not produce a spurious result. In cerebellar patients the largest order effect was an increase in 2.9° [ROPE:
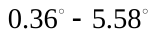
, 66% overlap with the ROPE] caused by the overlearning. We will address separately the possibility that order effects influenced our results, but what we see here is that these effects were small in general, although they could not be ruled out and were potentially as large as 1/8 to 1/4 of the size of our main effects. The full table can be seen online at https://github.com/Motor-Learning-Lab/paper-slowlearning.

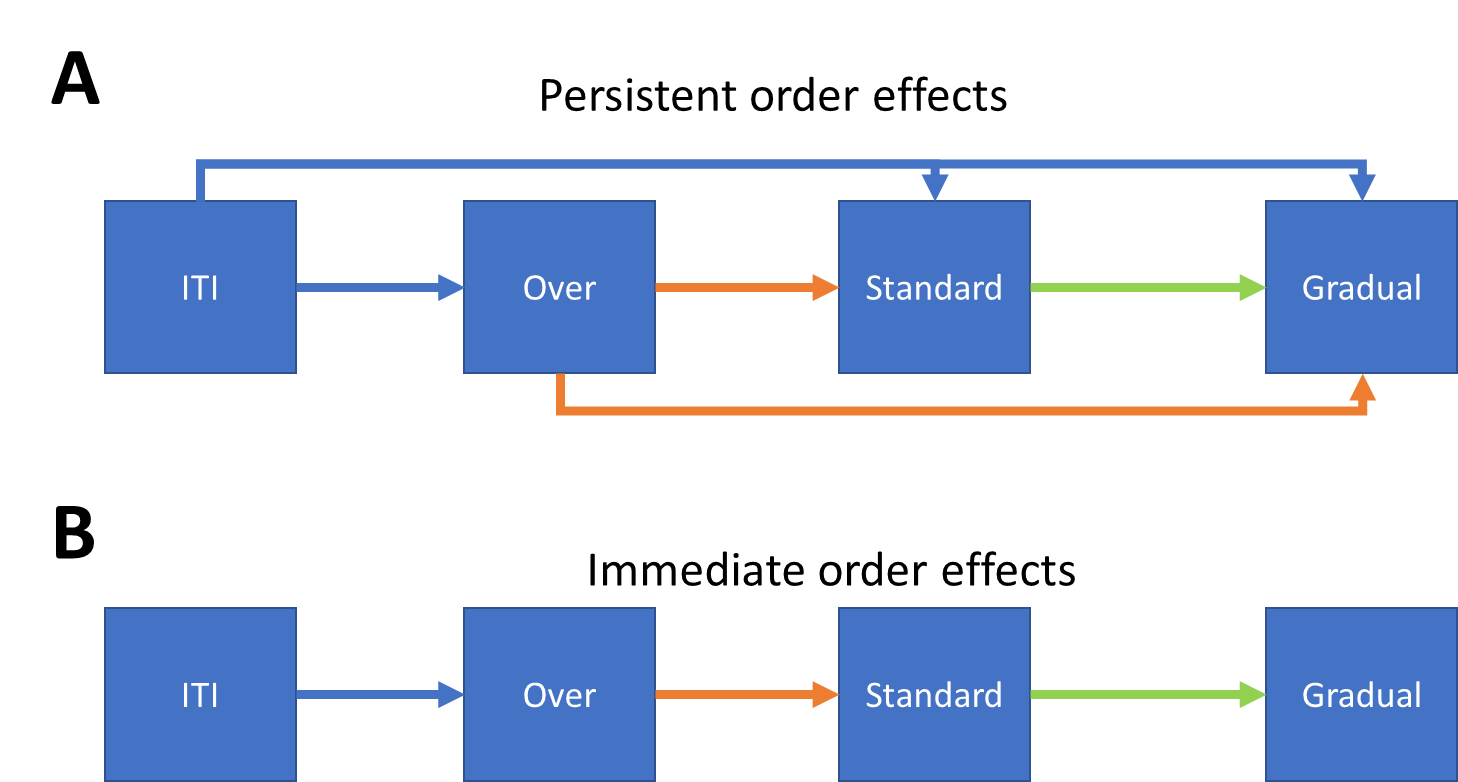

**Supplementary Figure 7**: Two models of the order effects. In the persistent effects model (A), the fact that one paradigm was done before the other is the same regardless of the number of intervening paradigms. In the immediate effects model (B), only the immediately preceding paradigm has an effect on the performance in the current paradigm.

We also re-examined how the two models of order effects influence the central results in the paper. **Supplementary Figure 8** shows the amount of spontaneous recovery in each of the three paradigms compared to the amount in the standard paradigm. As can be seen by comparing this figure to the results in **Main Text Figure 4**, accounting for order effects does not diminish the size of the effect we see and, if anything, strengthens it.

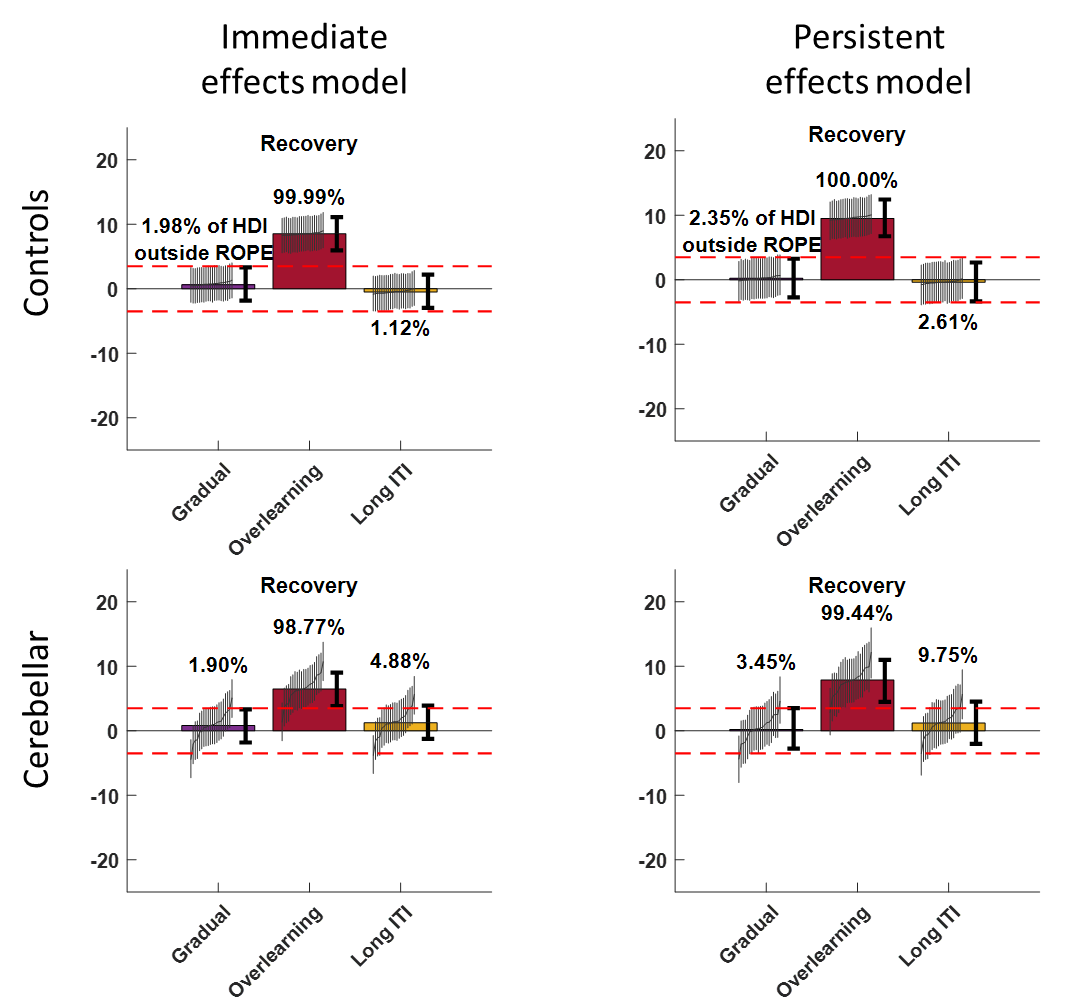

**Supplementary Figure 8**: The central result of the paper under different estimates of the order effect. The figure shows the difference between spontaneous recover in each paradigm and the standard paradigm in both control and cerebellar participants. This can be compared with the results without accounting for order effects as seen in Figure 4 of the main text.
